# Projections between the globus pallidus externa and cortex span motor and non-motor regions

**DOI:** 10.1101/2024.11.08.622712

**Authors:** E. A. Ferenczi, W. Wang, A. Biswas, T. Pottala, Y. Dong, A. K. Chan, M. A. Albanese, R. S. Sohur, T. Jia, K.J. Mastro, B. L. Sabatini

## Abstract

The globus pallidus externa (GPe) is a heterogenous nucleus of the basal ganglia, with intricate connections to other basal ganglia nuclei, as well as direct connections to the cortex. The anatomic, molecular and electrophysiologic properties of cortex-projecting pallidocortical neurons are not well characterized. Here we show that pallidocortical neurons project to diverse motor and non-motor cortical regions, are organized topographically in the GPe, and segregate into two distinct electrophysiological and molecular phenotypes. In addition, we find that the GPe receives direct synaptic input back from deep layers of diverse motor and non-motor cortical regions, some of which form reciprocal connections onto pallidocortical neurons. These results demonstrate the existence of a fast, closed-loop circuit between the GPe and the cortex which is ideally positioned to integrate information about behavioral goals, internal states, and environmental cues to rapidly modulate behavior.

## Introduction

The globus pallidus externa (GPe) is a central nucleus of the basal ganglia whose function has been implicated in diverse behaviors including regulation of movement and reward-related actions^1–4^. The GPe was canonically considered a “motor relay” within the indirect pathway of the basal ganglia, exerting an overall suppressive effect on motor output^5–8^. However, it is now apparent that the GPe plays a role in complex goal-directed behavior, mediated by a heterogeneous neural population that forms connections to a broad array of brain regions, both within and outside the basal ganglia^1–3,9,10^. Understanding how the GPe coordinates and transforms the flow of neural activity within the basal ganglia and to connected brain regions is critical for understanding the pathophysiology of disorders of basal ganglia circuits, such as Parkinson’s disease, dystonia, Tourette’s syndrome and others^1,11–21^.

*In vivo* electrophysiological recordings were first performed in the globus pallidus over 60 years ago^22–27^. These revealed that activity of pallidal neurons is correlated with features of movement^23–25,27,28^ and reward^26^. Two distinct electrophysiological cellular phenotypes were identified which differ in spontaneous action potential firing rate and bursting properties^23,25^. Since then, the superposition of rich anatomic, molecular, and electrophysiologic information has generated an updated understanding of the complex tapestry of cell types within the GPe^1,4,12,29–37^. The current working model subdivides the GPe neural population into two main subtypes. First, the canonical GABAergic “prototypic” neurons, which comprise approximately 70% of GPe neurons, express genetic markers such as *parvalbumin (Pvalb)* and *Nkx2-1*^32,38^, have high spontaneous firing rates, receive inputs from indirect pathway Type 2 dopamine receptor-expressing neurons from the striatum and send inhibitory outputs to traditional striatal indirect pathway output structures such as the substantia nigra pars reticulata (SNr)^4,23,29,30,32^. The second major subtype of GABAergic GPe neurons are referred to as “arkypallidal” neurons. These neurons express *FoxP2*, have lower baseline firing rate, and receive input from the direct pathway Type 1 dopamine receptor-expressing striatal neurons^39^ and send output primarily back to the striatum^31,32,38–40^. Within these two main subclasses, further subdivisions are classified according to the expression of variably overlapping profiles of molecular markers such as *Lhx6*^11,12,41^, *Npas1*^36,37,42,43^*, and Nkx2.1*^36,38^. Recently, single cell sequencing of the GPe generated an array of novel genetic markers that segregate the GPe into seven distinct molecular subclusters^44^, and detailed anatomic studies have identified rarer GPe neural populations that may be missed with bulk sequencing approaches^33,45–47^. In particular, a small neural population (∼5% of GPe neurons) sends a unique direct projection to the cortex and expresses markers for GABAergic and cholinergic neurotransmission^33,45–49^. The discovery of this pallidocortical neuron population reshaped our understanding of the GPe not as an intermediate relay station within the basal ganglia, but as an output nucleus with the ability to directly modulate cortical activity.

The ability of GPe to communicate rapidly and monosynaptically with the cortex raises critical questions about the role of pallidocortical circuits in integrating information about behavioral goals, internal states, and the external environment to modulate complex behaviors. However, it is important to first understand the detailed properties and organization of pallidocortical circuits. Previous work has shown that topographic heterogeneity of the GPe is layered over its molecular heterogeneity. The topographic organization of striatal inputs is maintained at the level of GPe and propagated to downstream structures, including the cortex^50,51^. However, the detailed topographic organization of pallidocortical neurons, the extent of cortical regions to which they project, and the nature of reciprocal topographic input from the cortex back to the GPe remains unknown.

The presence of pallidocortical and corticopallidal circuits is relevant for human disease, as white matter tracts connecting cortical to basal ganglia structures have become important targets for surgical intervention for neurologic and psychiatric disease. In humans, the “motor hyperdirect” pathway from the motor cortex directly to the STN is considered a key target for the therapeutic benefits of deep brain stimulation in Parkinson’s disease^52,53^. Other “hyperdirect” white matter bundles coursing from cortical regions such as the cingulate cortex to the basal ganglia have been postulated as DBS targets for other neuropsychiatric conditions such as obsessive compulsive disorder^54^. Understanding the detailed molecular, anatomic and physiological properties of circuits between the GPe and the cortex will enhance our ability to identify and optimize novel approaches and targets for neuromodulatory therapy.

Here we examined the anatomic and topographic distribution of pallidocortical neurons within GPe and correlated their electrophysiological properties with their cortical projection target. In addition, we examined the molecular markers that characterize anatomically defined pallidocortical neurons. Lastly, we examined if pallidocortical neurons receive direct monosynaptic input from the cortex in a “closed loop” manner, and if so, how this input differs from cortical input to other GPe cell types. Our findings indicate that pallidocortical neurons project to both motor and non-motor regions of cortex, and their topography is reflected in their cortical projection target. Pallidocortical neurons segregate into two distinct electrophysiological and molecular subtypes and receive direct monosynaptic excitatory input back from motor and non-motor regions of cortex, which is comparable in magnitude and latency to cortical input to other GPe neurons.

## Results

### Pallidocortical neurons encircle GPe with a topography that depends upon cortical projection target

Prior studies have suggested that pallidocortical neurons congregate at the medial and ventral borders of GPe^33,45–47^, but it is unknown whether they maintain the spatial topography of their cortical targets. To determine whether the spatial distribution of pallidocortical neurons varies according to their cortical projection target, we chose to sample four different cortical regions, including both sensorimotor regions – primary and secondary motor cortex (from now on referred to as “motor cortex”), primary somatosensory cortex (“sensory cortex”) – and higher order association areas – anterior cingulate cortex, prelimbic and infralimbic cortex (“frontal cortex”) and dorsal agranular insular cortex and gustatory (dysgranular and granular) insular cortex (“insular cortex”). We injected a retrograde fluorophore-encoding, G-deleted, non-pseudotyped rabies virus (RV) into each of these cortical regions to determine the spatial distribution of pallidocortical neuron cell bodies in GPe (Fig. 1a). This virus directly infects axons in cortex and expresses a fluorophore to fill the cell body and reveal its location. The injections were performed in *Drd2-GFP* transgenic mice, in which the pattern of GFP expression highlights the GPe^55^ (Table 1, Materials and Resources).

**Figure 1.**
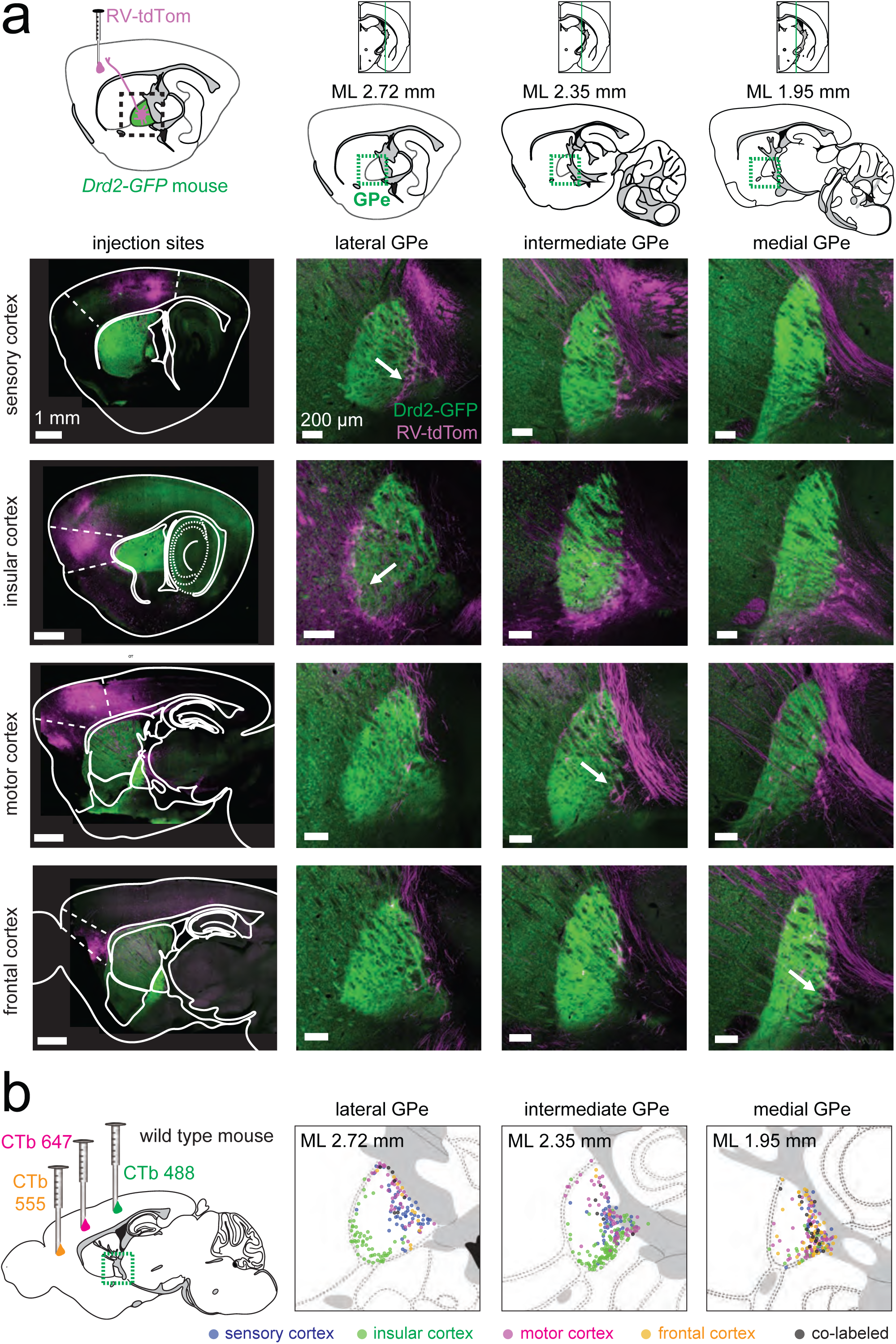
Pallidocortical neurons encircle GPe. **a,** *top,* Schematic of experimental approach: non-pseudotyped tdTom-expressing rabies virus (RV-tdTom) was injected into one of four cortical regions (sensory cortex, motor cortex, insular cortex, frontal cortex) in *Drd2*-*GFP* mice. *bottom,* Sagittal section fluorescence images showing the results for example mice. Injections sites are shown in the left panel. RV-infected cells (magenta) in GPe (green) are shown at lateral, intermediate and medial planes through GPe (arrows provide an example). GPe is highlighted by GFP fluorescence in indirect pathway (*Drd2*) axons (n=4 mice, 1 per cortical injection site). **b,** Cholera-toxin b (CTb) tagged with different fluorophores was injected into three different cortical regions in each mouse. The location of CTb-labeled cells in GPe annotated on sagittal section diagrams from the Allen Brain Atlas (n=563 cells across 6 mice, 3 cortical injections per mouse).

**Table 1.**
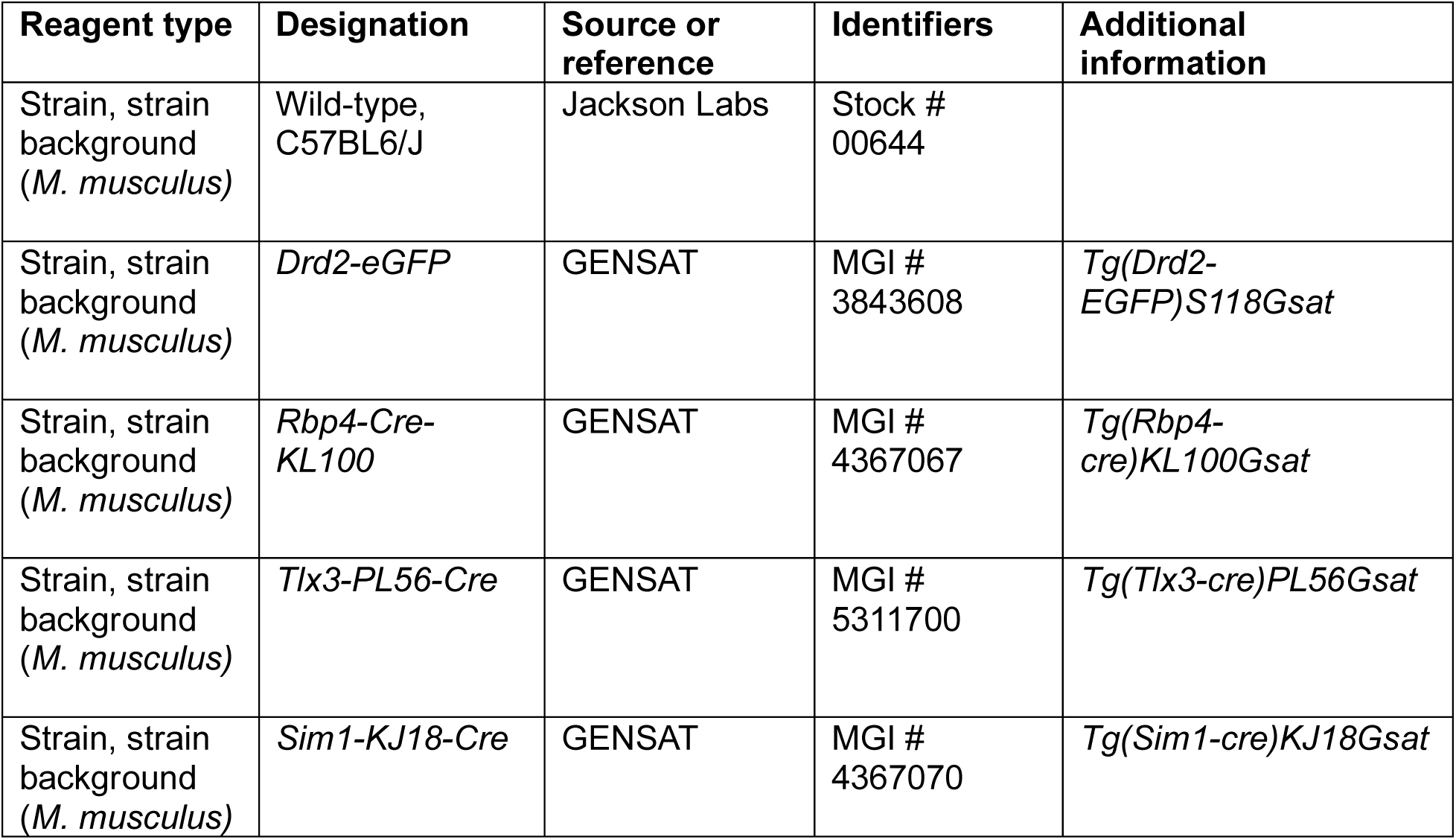
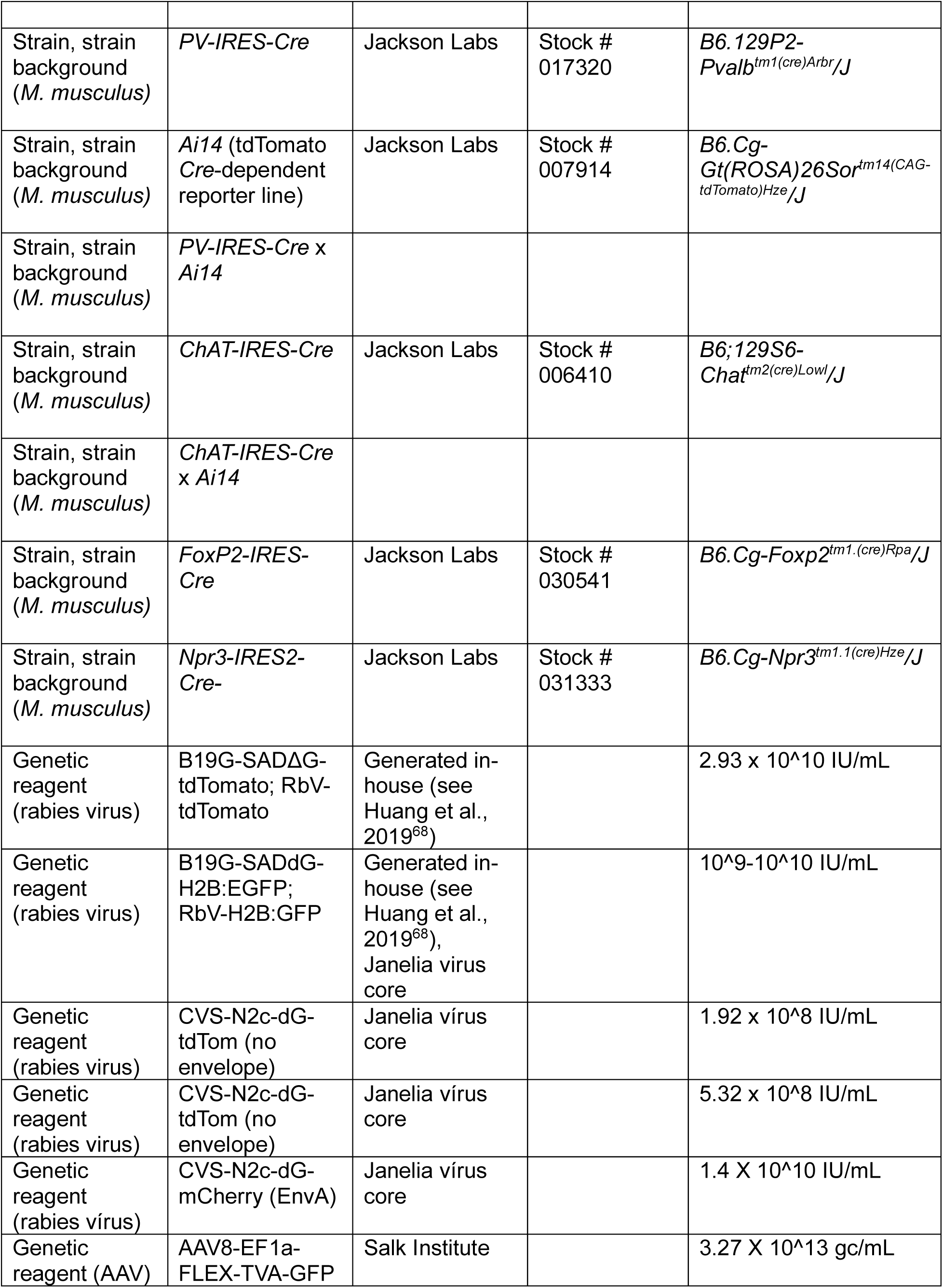
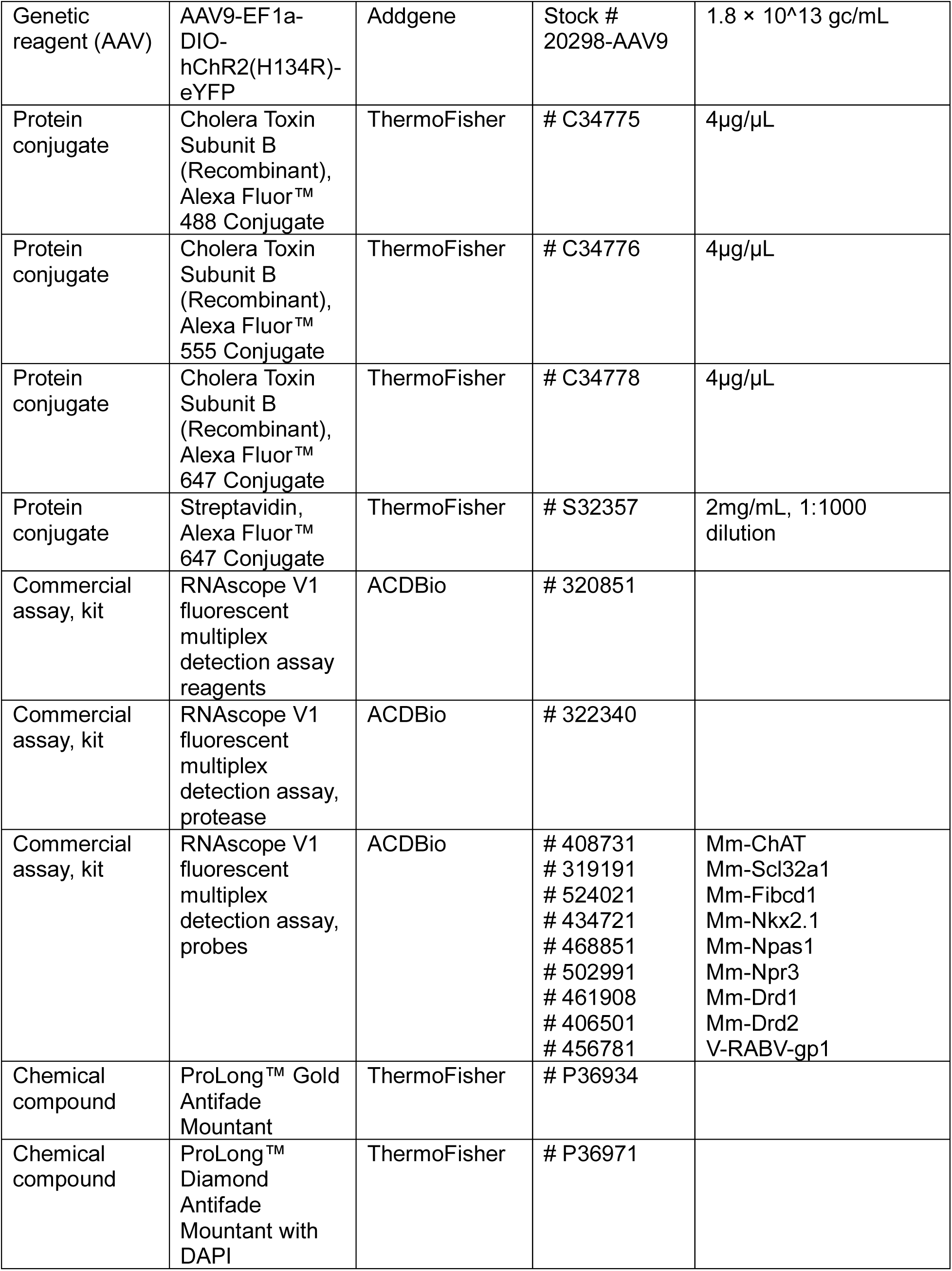
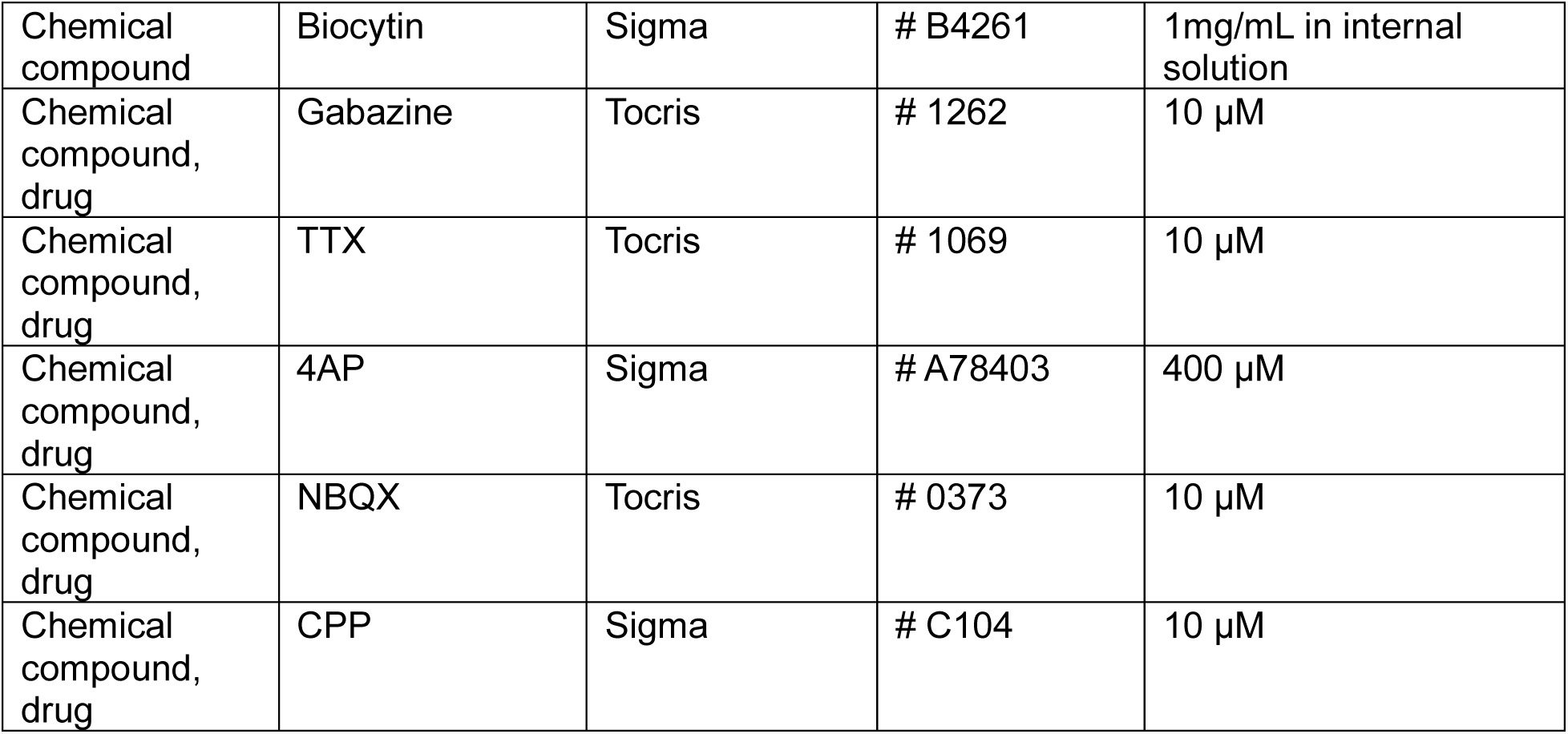
Materials and Resources.

We observed that pallidocortical neurons encircle the GPe in a ring-like manner and neurons that project to different cortical regions tend to align to different GPe borders. For example, insular- and sensory-cortex projecting neurons are numerous along the anterolateral border of GPe, whereas prefrontal cortex and motor cortex-projecting neurons are most concentrated at posteromedial borders (Fig. 1a). To verify that this spatial segregation was not a confound of viral tropism, we performed a similar experiment using a non-viral retrograde tracer, cholera toxin b (CTb) (Fig. 1b). Using CTb labeled with three different fluorophores, we assessed the topography of pallidocortical neurons projecting to three different cortical regions simultaneously and obtained similar results as with RV (Fig. 1b, Supp. Fig. 1c). Only a small proportion of neurons were co-labeled with more than one CTb fluorophore, suggesting that GPe neurons tend to project to specific cortical areas, rather than sending wide-reaching collaterals across multiple different cortical regions (Supp. Fig. 1c, d). Although under-labeling of pallidocortical neurons may be underestimate the fraction of cells that innervated multiple regions, a positive control with injection of two different CTb fluorophore labels into the same cortical location generated a high proportion of co-labeled pallidocortical neurons (Supp. Fig. 2a-d).

### Pallidocortical neurons fall into two distinct electrophysiological clusters that do not depend upon cortical projection target

The functional impact of synaptic input from GPe to the cortex^33,45^ will depend in part upon the intrinsic electrophysiological properties of pallidocortical neurons. Prior work has shown that *Chat+* and *Slc32a1+* (vesicular GABA transporter) GPe neurons have distinct electrophysiological phenotypes^45^; however, it is unknown if pallidocortical neurons segregate into distinct electrophysiological subtypes according to their cortical target. To examine this, we retrogradely labeled pallidocortical neurons according to cortical projection target with CTb and performed whole-cell electrophysiological current clamp recordings *ex vivo* from CTb-labeled neurons in the GPe in acute brain slices (Fig. 2a). These recordings revealed that pallidocortical neurons exhibit a wide variability in electrophysiological properties. Some neurons exhibit slow, regular spiking whereas others fast spiking behavior with adaptation, with varying degrees of a sag response to hyperpolarization (Fig. 2b-d). However, this variability in intrinsic properties across neurons did not clearly correspond to differences in cortical projection targets. Nevertheless, some differences exist between motor/sensory cortex-projecting neurons and insular cortex-projecting neurons in terms of their response to hyperpolarization (sag potential greater for insular cortex-projecting neurons) (Fig. 2c), and the interspike interval during a spike train (shorter for insular cortex-projecting neurons) (Fig. 2d). Principal component analysis (PCA) across a wide range of electrophysiological properties (Fig. 2e, Supp. Fig. 3b,c) revealed that when analyzed independently of cortical projection target, pallidocortical neurons cluster into two distinct phenotypes, predominantly driven by action potential firing properties (inter-spike interval, current step for first spike, and action potential acceleration), and membrane resistance (Fig. 2e, Supp. Fig 3c).

**Figure 2.**
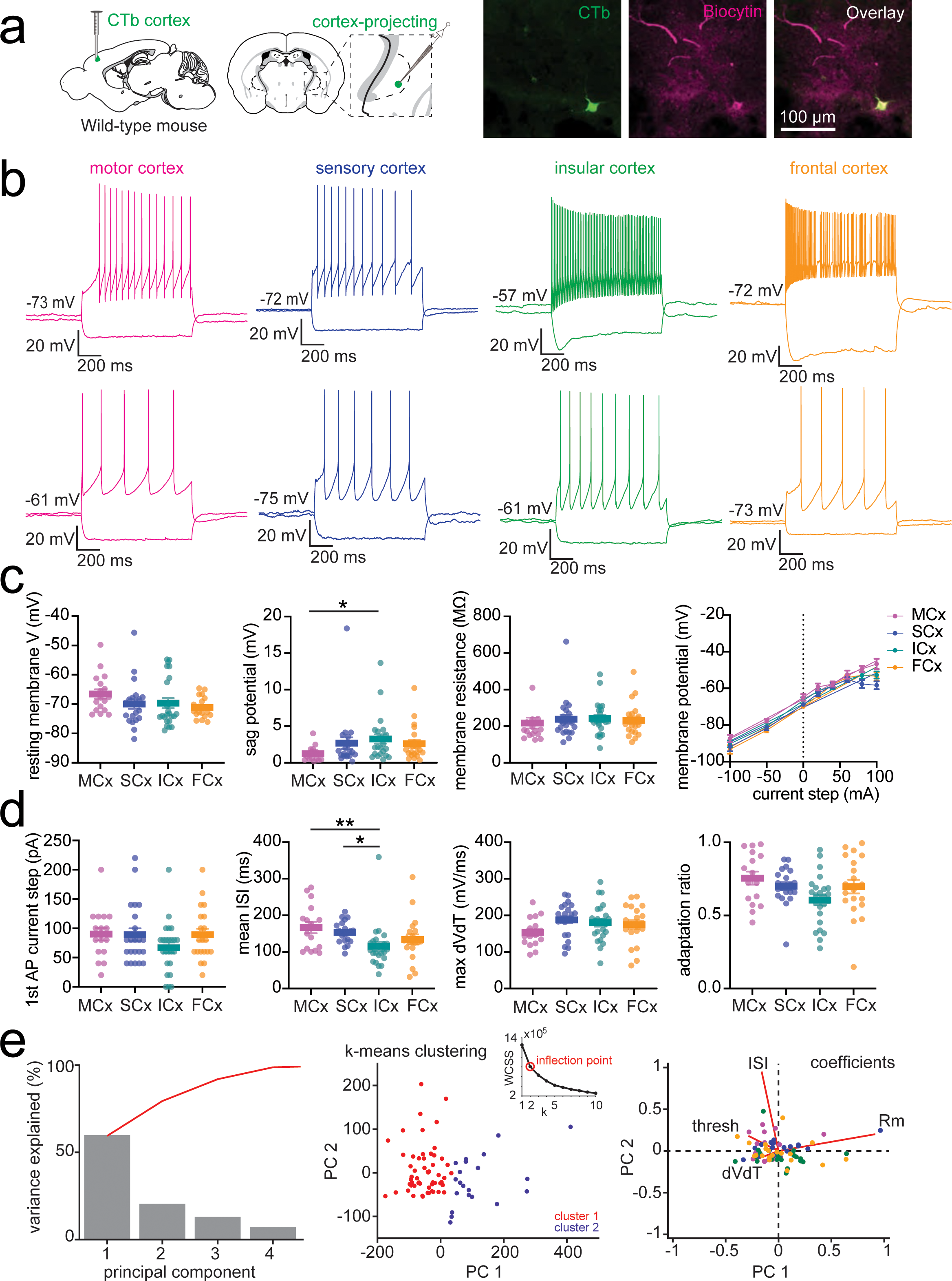
Pallidocortical neurons fall into two distinct electrophysiological clusters. **a,** *left,* Schematic of experimental approach: CTb was injected into one of four cortical regions – motor cortex (n=3 mice), sensory cortex (n=3 mice), insular cortex (n=3 mice), and frontal cortex (n=4 mice) – in wild-type mice to retrogradely label pallidocortical neurons in GPe for *in vitro* whole-cell recordings. *right,* Example image of a CTb-positive (green) and biocytin-positive (magenta) cell. Biocytin was introduced into the cell during the whole-cell recording for post-hoc identification. **b,** Example current clamp recordings from pallidocortical neurons projecting to each of the four cortical regions. A hyperpolarizing and a supra-threshold depolarizing response is shown for each cell. **c,** Quantification of passive properties of pallidocortical neurons according to cortical projection target: resting membrane potential (resting membrane V), sag potential (sag V), membrane resistance (Rm) and current-voltage relationship. Total number of recorded cells for each region: motor cortex: 17, sensory cortex: 22, insular cortex: 24, frontal cortex: 20. Mean and SEM are shown. There was a significant difference in sag potential between groups for sag potential (Kruskal-Wallis (K-W)=9.102, p=0.028, significant for motor cortex vs insular cortex p=0.0180 after multiple comparison correction). No significant differences between groups for other passive properties were identified. **d,** Quantification of active properties of pallidocortical neurons according to cortical projection target: current step required to generate a first action potential (1st AP current step), mean interspike interval (ISI), mean action potential rate of voltage change (mean max dVdT), mean adaptation ratio (first AP interval/last AP interval). Mean and SEM are shown. There was a significant difference between groups for mean ISI (Kruskal-Wallis (K-W)=16.65, p=0.0008, p=0.0213, significant for motor cortex vs insular cortex (p=0.0107) and sensory cortex vs insular cortex (p=0.0014), after multiple comparison corrections). No significant differences between groups after multiple comparison correction for other active properties. **e**, Principal component analysis (PCA) of electrophysiological properties (n=13 mice, 83 cells). *left,* Scree plot results. Percent variance in the data explained by the first four principal components. *middle,* K-means clustering for k=2, *inset:* Elbow plot of Within-Cluster Sum of Squares (WCSS) for different values of k. The maximum of the ratio of the second derivative/first derivative was used to identify the optimal value for k (the inflection point of the curve). Data points are color-coded by cluster assignment. The plot illustrates the distribution of data across clusters. *right*, Biplot of the first two principal components (PC1 and PC2) showing variable coefficients and scores. Points represent the scores for each cell on PC1 and PC2 and are colored according to cortical projection target. Red lines indicate the direction and magnitude of variable coefficients, the line length is proportional to the contribution of each variable to the principal components.

### There are two main molecular phenotypes of pallidocortical neurons

Since intrinsic electrophysiological variation between pallidocortical neurons was not primarily determined by cortical projection target, we hypothesized that these differences result from variation in the molecular identity of pallidocortical neurons. To evaluate whether there are distinct molecular identities of pallidocortical neurons, we performed *in situ* mRNA hybridization on brain slices containing GPe, in which pallidocortical neurons were labeled using the retrograde non-pseudotyped RV tracer (Fig. 3). Based on prior literature we expected pallidocortical neurons to express markers for cholinergic and GABAergic neurotransmission^33,45^ and indeed confirmed that *Slc32a1 and Chat* mRNA co-localized with rabies virus mRNA (*gp1* gene) (Fig. 3a-c). 64% of all RV-infected cells expressed at least one marker, and of these, a small proportion (11.5%) co-expressed *Chat* and *Slc32a1* (17% *of* RV-infected *Chat+* neurons expressed *Slc32a1*, and 24% of RV-infected *Slc32a1+* neurons expressed *Chat*). Since *Slc32a1* is expressed by many GPe neurons, we examined whether other candidate mRNA markers for GPe neurons - *Npas1*^36,42,43^, *Nkx2-1*^36^, *Fibcd1*^44^, and *Npr3*^36,44^ - could specifically identify non-cholinergic pallidocortical neurons. All these markers were expressed by pallidocortical neurons to some degree, however only *Npr3* and *Nkx2-1* clearly demarcated pallidocortical neurons into two distinct molecular subtypes – *Npr3+/ Slc32a1+* neurons, and *Nkx2-1+/Chat+* neurons (Fig. 3d-g).

**Figure 3.**
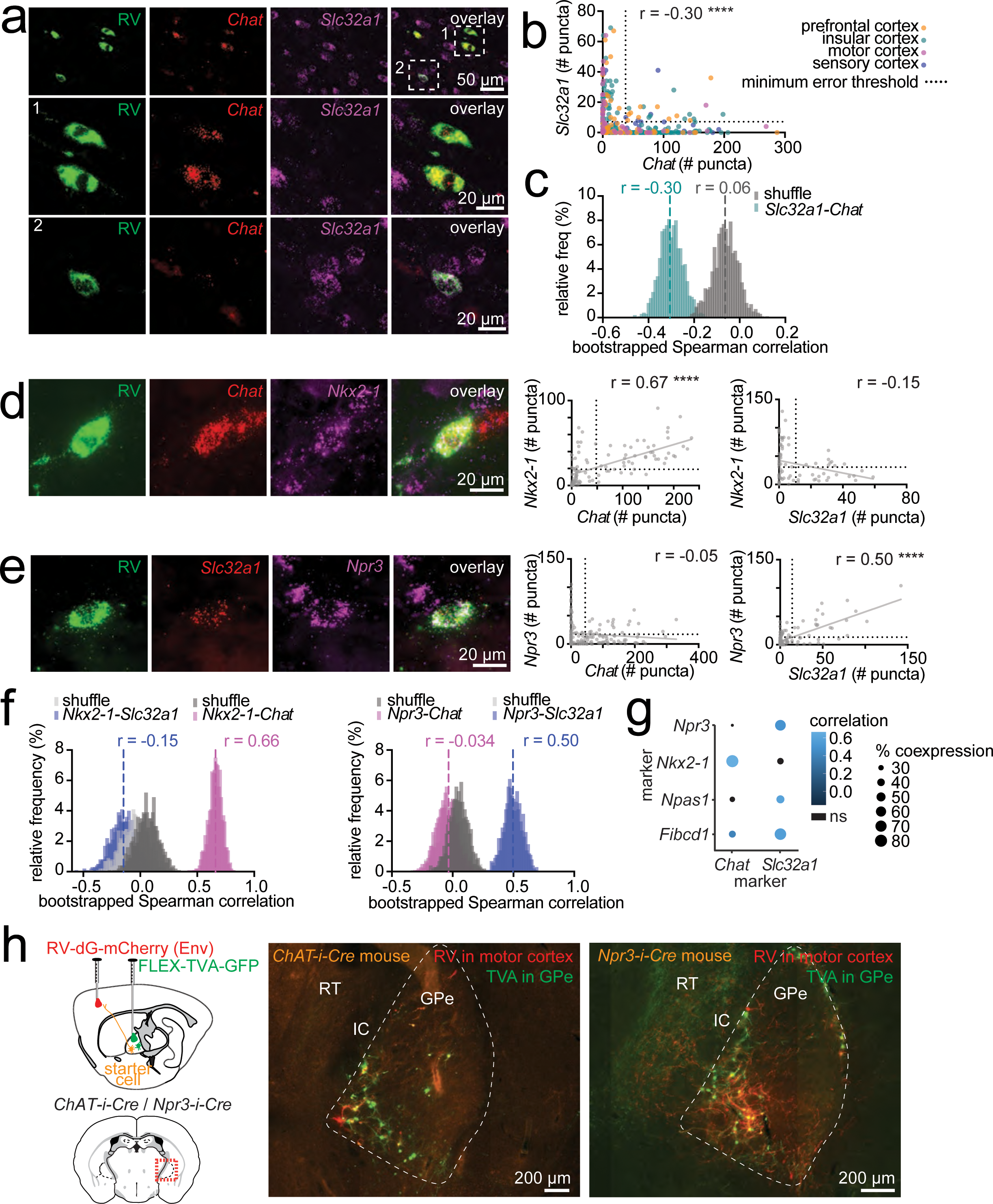
Two molecular classes of pallidocortical neurons. **a,** Rabies virus was used to retrogradely label pallidocortical neurons. *top row,* Example images of mRNA fluorescent *in situ* hybridization (mRNA-FISH), with labeling of rabies virus *gp1* (*RV*) and markers for cholinergic (*Chat* – choline acetyl transferase) and GABAergic (*Slc32a1* – vesicular GABA transporter) neurotransmission. *middle row,* enlarged image of *Chat*-positive/*Slc32a1*-negative RV infected neuron. *bottom row,* enlarged image of *Chat*-negative/*Slc32a1*-positive RV infected neuron. **b,** Quantification of mRNA puncta numbers for *Chat* and *Slc32a1* in RV-infected pallidocortical neurons. Each dot represents a different neuron, colored according to its cortical projection target. The minimum error threshold classifies neurons into expressing or non-expressing for each marker. Spearman rho (r) correlation coefficient for number of *Chat* puncta versus *Scl32a* puncta: -0.302 (p<0.0001, n=4 mice, 326 cells) **c,** Bootstrap analysis for Spearman correlation coefficients for non-shuffled and shuffled data: mean non-shuffled r: -0.3, mean shuffled r: -0.06 (standard deviation: 0.056, number of bootstraps=1000). **d,** Example images of mRNA-FISH with labeling of *RV*, *Chat* and *Nkx2-1*. Quantification of mRNA puncta for *Nkx2-1* vs *Chat* (Spearman rho: 0.67 (p < 0.0001, n=1 mouse, 106 cells) and *Nkx2-1* vs *Slc32a1* (Spearman rho: -0.15, p=0.2, n=1 mouse, 75 cells). **e**, Example images of mRNA-FISH with labeling of *RV*, *Slc32a1* and *Npr3*. Quantification of mRNA puncta for *Npr3* vs *Chat* (Spearman rho: -0.05 (p=0.58, n=1 mouse, 159 cells) and *Npr3* vs *Slc32a1* (Spearman rho: 0.50, p < 0.0001, n=1 mouse, 126 cells). **f,** Bootstrap analysis for Spearman correlation coefficients for non-shuffled and shuffled data for *Nkx2-1* vs *Chat* and *Nkx2-1* vs *Slc32a1; Npr3* vs *Chat* and *Npr3* vs *Slc32a1.* Mean correlation coefficients for non-shuffled data are shown, along with distribution of shuffled and non-shuffled bootstrapped data. Distributions of non-shuffled data for *Nkx2-1* vs *Chat* and *Npr3* vs *Slc32a1* are separated from their respective shuffled distributions by more than 2 standard deviations. **g,** Dot plot summary for correlation coefficients and percent co-expression between different GPe markers in pallidocortical neurons. Non-significant correlation coefficients are indicated in black. **h.** Confirmation that *Chat* and *Npr3* can be used to label pallidocortical neurons in GPe. Experimental approach: *Cre*-dependent TVA-GFP helper virus was injected into GPe of *Chat-i-Cre* or *Npr3-i-Cre* mice, followed 2 weeks later by injection of G-deleted, EnvA-pseudotyped rabies virus (CVS-N2c-dG-mCherry (EnvA)) into either motor cortex or insular cortex. Example images of TVA-expressing (green) and rabies expressing pallidocortical neurons (red/yellow) in GPe in *Chat-i-Cre* (left image) and *Npr3-i-Cre* (right image).

To confirm that our mRNA *in situ* hybridization experiments reliably identified genetic markers that selectively label pallidocortical neurons, we used pseudotyped RV to retrogradely label pallidocortical neurons in a genetically specified manner. Pseudotyped RV contains an envelope protein that requires a helper virus (avian tumor virus receptor A, TVA) to infect axonal terminals. TVA expression can be made *Cre*-recombinase dependent to restrict rabies infection to only the subset of neurons that express *Cre*. We injected a *Cre*-dependent TVA helper virus into the GPe of *Npr3-i-Cre* (*Npr3-IRES-Cre*) or *ChAT-i-Cre (ChAT-IRES-Cre)* transgenic mice (Table 1, Materials and Resources), to drive TVA expression selectively in either Npr3-positive (Npr3+) or ChAT-positive (ChAT+) neurons respectively. Two weeks later, we injected EnvA pseudotyped, G- deleted RV into either motor or insular cortex of the same mice (Fig. 3h, Supp. Fig. 5c). This resulted in strong rabies virus labeling in GPe neurons in transgenic mice (Fig. 3h) but not wild type control mice (Supp. Fig. 5c), confirming the existence of cortically-projecting Npr3+ and ChAT+ cells in GPe. This approach also allowed us to visualize axonal collaterals of Npr3+ and ChAT+ neurons in other brain regions, including Npr3+ axons in the reticular nucleus of the thalamus (RT), consistent with prior reports^35^. In contrast, ChAT+ axons were not seen in the RT but were seen in the amygdala (Supp. Fig. 5b). This suggests that pallidocortical neurons project not only to cortex but also to other subcortical targets, and the distribution of these may differ according to their molecular identity.

### Pallidal neurons receive monosynaptic glutamatergic input from deep layers of motor and non-motor cortical regions

The GPe receives direct synaptic input from primary and secondary motor cortex^36,56^ and but the extent to which other cortical regions also synapse on GPe neurons and which neurons receive this input remains unknown. Having confirmed that pallidocortical neurons project to both sensorimotor and higher order association areas of cortex, we examined if GPe neurons receive input directly back from non-motor, and higher order cortical regions. To assess this anatomically, we injected a non-pseudotyped RV encoding a fluorophore into GPe to generate a cortex-wide map of neurons whose axon terminals innervate the GPe (Fig. 4a, Supp. Fig. 6, 7). As a control, we simultaneously injected a similar RV expressing a different fluorophore into the striatum and compared the distribution and pattern of inputs between the two basal ganglia regions. We observed that inputs to GPe and striatum arose predominantly from neurons on the side of the brain ipsilateral to the injection site (Fig. 4b). Furthermore, the cortex was the second most common site for input to GPe, after the striatum (Fig. 4c, Supp. Fig. 7a,c). The mean total number of cortical cells projecting to GPe was 26,616 and mean total number of cortical cells projecting to striatum was 25,455. When analyzing according to cortical layer, we found that the majority of GPe input from cortex arises from deep cortical layers, in particular, layer V, whereas input to the striatum arises from both deep and more superficial cortical layers, layers II and III (Fig. 4d). Cortical input to GPe arises from a large swath of cortical regions. Sensorimotor cortex provides the largest input, followed closely by higher order association areas such as the agranular insular cortex (Supp. Fig 6, 7b-d).

**Figure 4.**
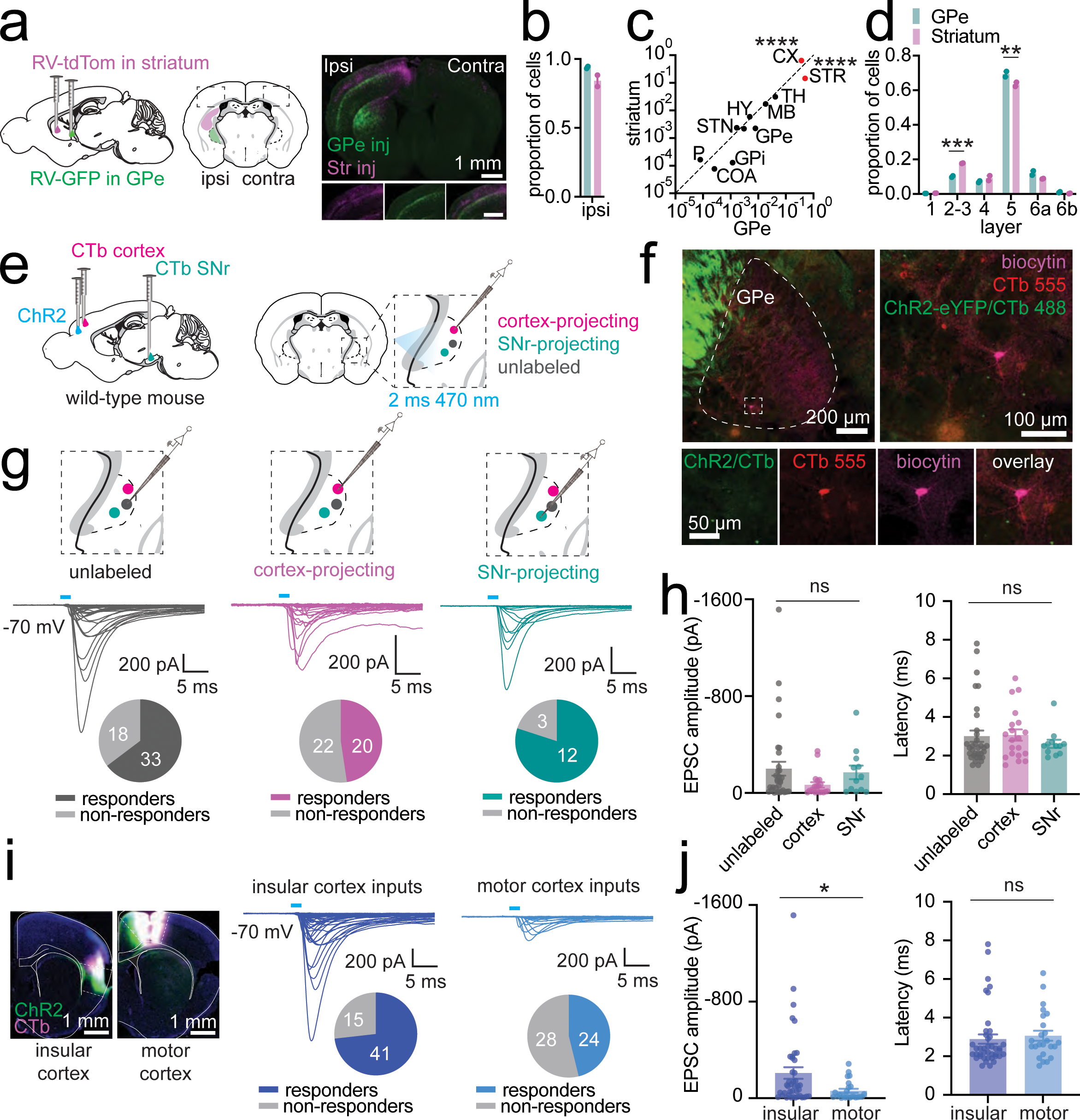
Cortical input to GPe neurons. **a,** Anatomic tracing of cortical inputs to GPe. *left,* Schematic of experimental approach: non-pseudotyped G-deleted RVs encoding either GFP or tdTomato was injected into GPe (CVS-N2c-dG-GFP) and striatum (CVS-N2c-dG-tdTomato) of the right hemisphere of wild type mice (n=3). *right,* Example images of cortex ipsilateral and contralateral to the injection site. *inset*, zoom of cortical region. **b,** Proportion of all cells innervating GPe and striatum ipsilateral to the injection site (n=2 mice, mean (+/- SEM) total cells for GPe=72400 (13563), mean (+/- SEM) total cells for striatum=39834 (1754). **c,** Proportion of ispsilateral cells innervating GPe vs striatum, stratified by broad brain regions (n=2 mice). There was a significant difference between GPe and striatum for cortical input (p<0.0001) and striatal input (p<0.0001) after multiple comparison correction (two-way ANOVA with Šídák’s multiple comparison test). *Key,* STR: striatum, CX: cortex, TH: thalamus, MB: midbrain, HY: hypothalamus, STN: subthalamic nucleus, GPe: globus pallidus externa, GPi: globus pallidus interna, HPF: hippocampal formation, COA: cortical amygdala area, P: pons. **d,** Proportion of all ipsilateral cortical cells innervating GPe vs striatum, stratified by cortical layer (n=2 mice). There was a significant difference between GPe and striatum for layers 2-3 (p=0.0001), and layer 5 (p=0.0014) after multiple comparison correction (two-way ANOVA with Šídák’s multiple comparison test). **e,** Schematic for experimental electrophysiological verification of cortical innervation to GPe in wild type mice: CTb was injected into cortex and SNr to retrogradely label GPe neurons innervating cortex or SNr respectively. ChR2 was injected into cortex at the same site as CTb. Whole-cell voltage clamp recordings were obtained from CTb labeled or unlabeled GPe neurons and used to record synaptic currents evoked by stimulation of cortical axons with a single 2 ms blue light pulses. **f**, Example histological image of an acute brain slice used for electrophysiological analyses showing a biocytin and CTb-labeled cortex-projecting neuron analyzed by whole-cell recording. **g**, Synaptic currents evoked by ChR2 stimulation of cortical axons recording in three different GPe neuron types: unlabeled (n=51 cells, 14 mice), cortex-projecting (pallidocortical, n=42 cells, 15 mice), and SNr-projecting (pallidonigral, n=15 cells, 5 mice). The proportions of responding vs non-responding cells are shown below the traces. **h,** Summary of EPSC amplitude and onset latency for all responding neurons. Mean and SEM are shown. There was no significant difference between neuron types for EPSC amplitude (Kruskal-Wallis test K-W=3.914, p=0.1413) or latency (K-W=0.6003, p=0.7407). **i,** Example histology of ChR2 and CTb injections in insular cortex and motor cortex are shown. Synaptic currents evoked by ChR2 stimulation of axons from insular cortex (n=56 cells, 20 mice) and motor cortex (n=52 cells, 14 mice) to all GPe neuron types. **j**, Summary of EPSC amplitude and onset latency for all responding neurons. Mean and SEM are shown. EPSCs evoked by insular cortex stimulation to GPe neurons were significantly larger those evoked by motor cortex inputs (two-tailed Mann-Whitney U test, U=312, p=0.0139). There was no significant difference between onset latency (two-tailed Mann-Whitney U test, U=390, p=0.1670).

We examined if these anatomical observations indicate the existence of functional synapses and whether the strength of cortical synaptic input varies according to the projection target of different GPe neurons. Based on prior studies^56^, we hypothesized that the “traditional” prototypic output neurons of the GPe – for example, those projecting to the substantia nigra pars reticulata (SNr) – may receive weaker synaptic input from cortex compared to other GPe neuron types. To test this, we injected CTb labeled with one fluorophore into cortex and CTb labeled with a different fluorophore into the SNr of wild-type mice. In addition, we injected the optogenetic neural activator, channelrhodopsin (ChR2), into either motor or insular cortex (Fig. 4d,f) to activate cortical axonal terminals in the GPe with blue light in acute brain slices prepared 3-8 weeks after injection (mean=4.5 weeks, std = 1.2 weeks).

While collecting this data, we observed a proportion of GPe neurons with unusually fast excitatory currents evoked by pulses of blue light, with a latency of less than 1 ms from the onset of the blue light pulse (Supp. Fig. 8a). To determine whether these could be physiological, we attempted to block these currents using pharmacologic blockers of neuronal spiking (TTX 10 uM) and excitatory glutamatergic synaptic transmission (NBQX, CPP 10 µM), and found that they were not pharmacologically suppressible, consistent with direct activation of optogenetic photocurrents (Supp. Fig. 8a). Most neurons with photocurrents were pallidocortical neurons (those labeled with CTb injected into cortex), consistent with retrograde expression of ChR2 from cortex to GPe, even in the absence of clear ChR2-associated YFP expression in GPe. These short-latency currents were excluded.

After exclusion of these photocurrents, we were surprised to find that cortical-mediated synaptic currents were evoked in many neurons in GPe, including those projecting to SNr, as well as randomly selected neurons not labeled with any fluorophore (“unlabeled”) (Fig 4g, h). Importantly, we also identified synaptic input to cortex-projecting neurons, confirming the existence of a closed loop circuit in which pallidocortical neurons that project to the cortex receive direct excitatory input back from the cortex. The amplitude of excitatory synaptic currents did not significantly differ between these groups (Fig. 4h). However, the insular cortex synaptic inputs to GPe were more numerous and greater in amplitude than those from motor cortex (Fig. 4i, j).

### Pallidocortical neurons receive closed loop synaptic input from specific cortical layers

To eliminate off-target retrograde expression of ChR2 and examine the contribution of different cortical neurons to pallidal synapses, we repeated the same experiments in three different transgenic mouse lines to specifically target deep-layer cortical projection neurons. In an *Rbp4-Cre* mouse^55,57^ (Table 1, Materials and Resources), which targets layer 5 projection neurons, we again confirmed the presence of excitatory synaptic input to both unlabeled and pallidocortical neurons in GPe (Fig. 5a-d). With this approach, we no longer observed any fast photocurrents (< 1 ms from light pulse onset). We confirmed that the evoked currents were monosynaptic as they could be evoked in the presence of action potential blockade (TTX 10 uM) together with the potassium channel antagonist, 4AP (400 uM), which recovers synaptic vesicle release in the absence of action potential firing^58^ (Fig. 5c,d). Similarly, the currents were suppressed by blockers of glutamatergic neurotransmission (NBQX, CPP 10 µM), confirming that they are mediated by the excitatory neurotransmitter, glutamate. Since Rbp4-positive cortical neurons include both intra-telencephalic (IT) projection neurons and pyramidal tract (PT) projection neurons, we narrowed down the source of cortical input to GPe by repeating the experiments in either *Tlx3-PL56-Cre* mice^55,57^ to target IT neurons or *Sim1-KJ18-Cre* mice to target PT neurons^55,57^ (Fig. 5e, f) (Table 1, Materials and Resources). As a positive control for variability in ChR2 expression across mice, we also recorded from striatal neurons in the same brain slices and found that whereas *PT*-inputs to striatum were robust, they were small and infrequent to GPe neurons. In contrast, *IT*-inputs to striatum and GPe were of similar magnitude (Fig. 5e-f).

**Figure 5.**
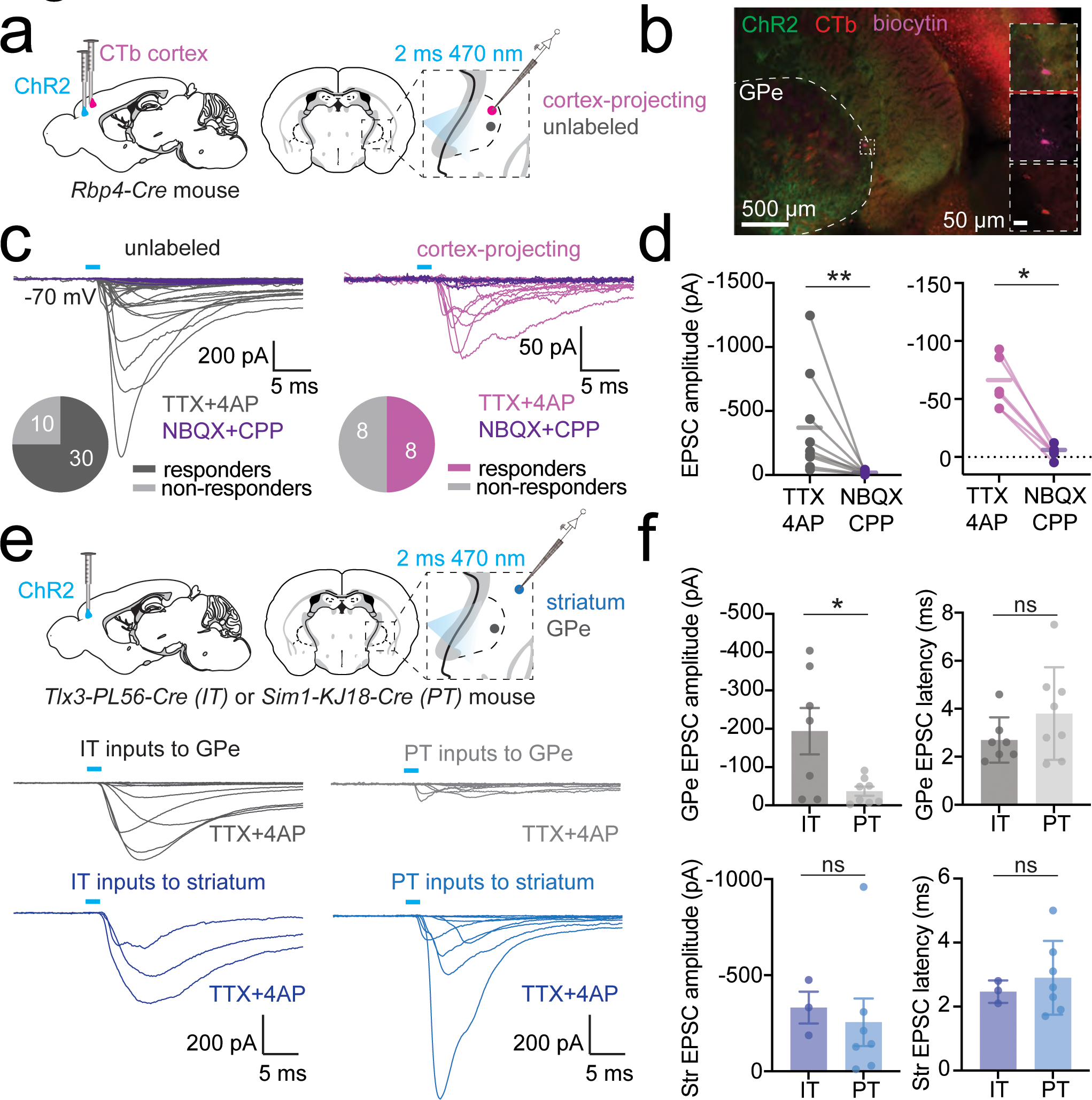
Cell-type specific cortical input to GPe neurons. **a,** Electrophysiological verification of cortical inputs to GPe in *Rbp4-Cre* mice. Schematic of experimental approach: CTb was injected into cortex to retrogradely label cortex-projecting GPe neurons. ChR2 was injected into cortex at the same site as CTb. CTb-labeled or unlabeled neurons were whole-cell voltage clamped in GPe, and single 2 ms 470 nm light pulses were used to stimulate *Rbp4^+^* cortical axons. Recordings were made in the presence of TTX and 4AP to isolate monosynaptic inputs, and in the absence or presence of blockers of glutamatergic synaptic transmission. **b**, Example histological image of an acute brain slice used for electrophysiological slice showing a biocytin-filled, CTb-labeled cortex-projecting neuron. **c**, Synaptic currents evoked by ChR2 stimulation of Rbp4+ cortical inputs for unlabeled (n=40 cells, 11 mice) and cortex-projecting (pallidocortical, n=16 cells, 8 mice) cells. Responses in the presence of glutamatergic synaptic blockers are shown (purple traces) for a subset of cells. The proportion of responding vs non-responding cells in the absence of synaptic blockers are shown below the traces. **d,** Summary of EPSC amplitude for neurons in the absence and presence of glutamatergic synaptic blockers. Mean and SEM are shown. Responses were significantly reduced by synaptic blockers for both unlabeled and cortex-projecting neurons. Unlabeled cells: n=9 cells, 9 mice: Wilcoxin matched-pairs signed rank test W=45, p=0.0039; cortex-projecting cells: n=6 cells, 3 mice: W=21, p=0.0312. **e,** Schematic of experimental approach: ChR2 was injected into the cortex of either *Tlx3-PL56-Cre* mice (to target intratelencephalic, IT, neurons) or *Sim1-KJ18-Cre* mice (to target pyramidal tract, PT, neurons) and unlabeled neurons were whole-cell voltage clamped in GPe and striatum. 2 ms blue light pulses were used to stimulate IT or PT cortical axons in GPe, in the presence of TTX and 4AP to isolate monosynaptic inputs. Synaptic currents evoked by ChR2 stimulation of IT axons in GPe (n=15 cells, 2 mice) and striatum (n=3 cells, 2 mice), and PT axons in GPe (n=22 cells, 2 mice) and striatum (n=9 cells, 2 mice). **f,** Summary of EPSC amplitudes and onset latency. Mean and SEM are shown. There was a significant difference in EPSC amplitude for IT vs PT input to GPe neurons (Mann Whitney test: U=9, p=0.0289) but not for striatal neurons (U=5, p=0.2667). There was no difference in onset latency between IT and PT input for either GPe neuron class.

### The functional impact of cortical excitatory input to GPe is determined by the molecular and electrophysiological properties of different GPe neurons

To assess the functional impact of excitatory cortical input on spiking activity of GPe neurons, we recorded from molecularly defined neuron types in GPe during optogenetic stimulation of cortical axons from either motor cortex or insular cortex. We used transgenic mouse lines that permit conditional *Cre-*dependent fluorophore labeling of molecularly defined neurons (Table 1, Materials and Resources). The neuron subtypes targeted were canonical GPe parvalbumin-expressing “prototypical” neurons (Pvalb*+,* labeled using *Pvalb-i-Cre* transgenic mice crossed to a tdTomato transgenic reporter line, *Ai14*), FoxP*2-*expressing “arkypallidal” neurons^40^ (FoxP2*+*, targeted using virally-mediated *Cre-*dependent fluorophore expression in *FoxP2-i-Cre* mice) and ChAT*-* expressing neurons (ChAT*+, ChAT-i-Cre* x *Ai14* transgenic cross) (Fig. 6). We recorded excitatory synaptic currents in all neuron types and found no significant difference in amplitude of EPSCs to the different cell types (Fig. 6b). However, when assessing the membrane potential response to this synaptic input in current clamp mode in the same neurons, we found that this input evoked spikes more frequently in ChAT+ neurons than in Pvalb+ neurons (Fig. 6c,d), which may in part be due to differences in intrinsic membrane properties such as resting membrane potential (Supp. Fig. 10b). Although light pulses did not evoke spikes in Pvalb*+* neurons that were not spiking spontaneously, in a subset of spontaneously firing Pvalb*+* neurons, we found that light pulses reduced the interspike interval immediately following the light pulse, effectively resetting the phase of firing (Fig. 6f). To examine if an unbiased analysis of intrinsic electrophysiological properties segregates the data from all three cell-types into clusters according to molecular phenotype, we used PCA. PCA across a range of intrinsic electrophysiological properties revealed three distinct clusters, two of which segregated broadly into Pvalb*+-*like or ChAT+-like identities, and a third cluster that contained a mixture of all three cell-types (Supp. Fig. 12b, c). Key electrophysiological features that distinguished the clusters were membrane resistance, mean interspike interval, current step to generate a first action potential and the maximum firing frequency (Supp. Fig. 11, 12d).

**Figure 6.**
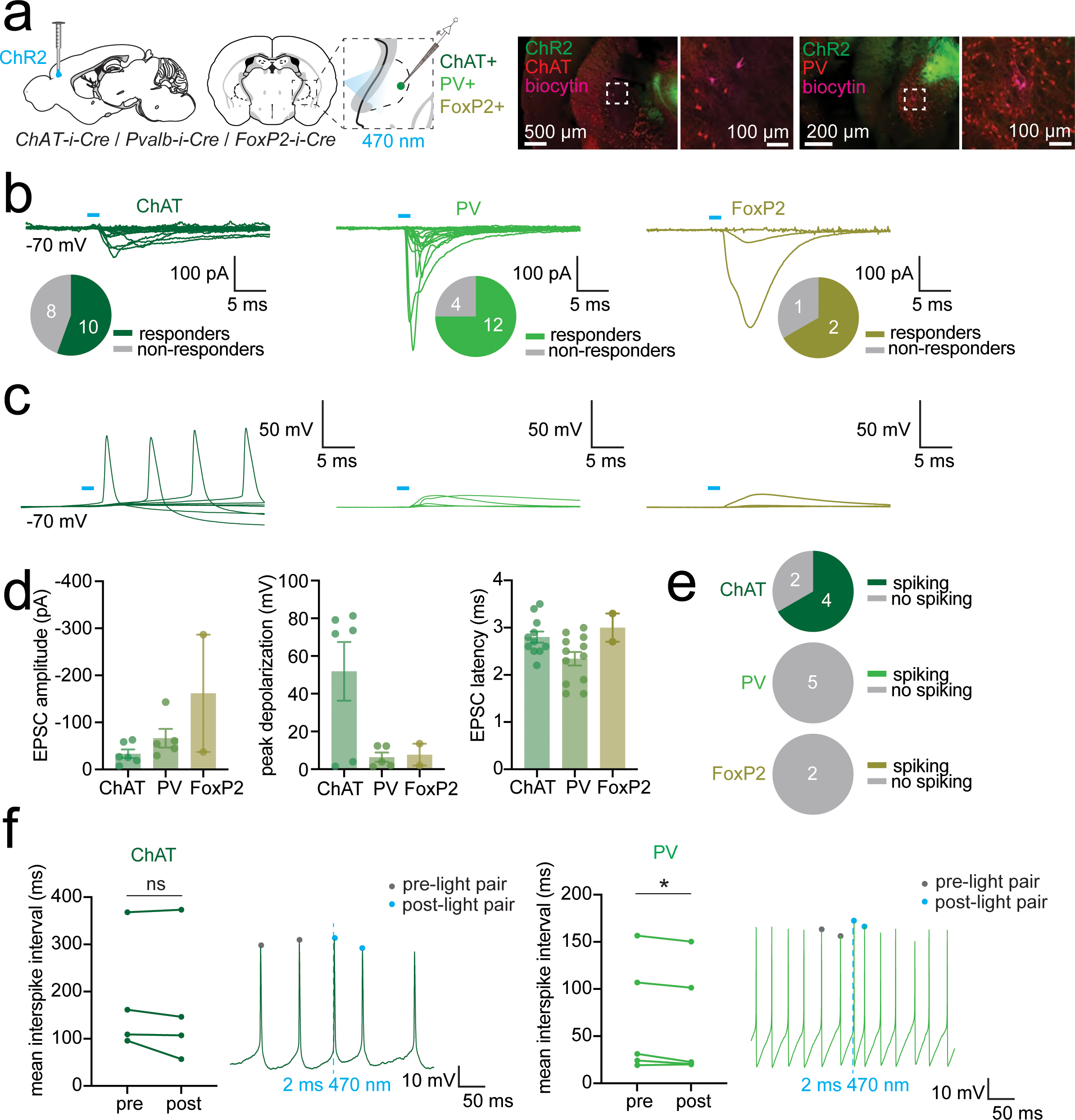
Cortical input to specific cell types in GPe. **a,** Schematic of experimental approach: ChR2 was injected into motor or insular cortex of *ChAT-i-Cre* x Ai14 (tdTom reporter), *Pvalb-i-Cre* x Ai14 (tdTom reporter), or *FoxP2-i-Cre* mice. An AAV carrying a *Cre*-dependent fluorophore was injected into GPe of *FoxP2-i-Cre* mice. tdTom-labeled (ChAT+ or Pvalb+) or fluorophore-labeled (FoxP2+) neurons were voltage-clamped in GPe. 2ms pulses of 470 nm light were delivered to stimulate cortical inputs to GPe. Example histological images are shown, biocytin indicates a recorded neuron. **b,** Synaptic currents evoked by ChR2 stimulation of cortical inputs to ChAT+, Pvalb+, and FoxP2+ neurons are shown. Proportion of responding vs non-responding cells are shown below traces. Number of mice/cells: ChAT n=2 mice, 18 cells; PV n=2 mice, 16 cells; FoxP2 n=2 mice, 3 cells. **c,** Synaptic potentials evoked by ChR2 stimulation of cortical inputs to ChAT+, Pvalb+, and FoxP2+ neurons. Number of mice/cells: ChAT n=2 mice, 6 cells, PV n=2 mice, 5 cells, FoxP2 n=2 mice, 2 cells. **d,** Summary of EPSC amplitude and peak depolarization evoked by ChR2 stimulation for all neurons for which both voltage- and current-clamp recordings were obtained. Number of mice/cells: ChAT n=2 mice, 6 cells, PV n=2 mice, 5 cells, FoxP2 n=2 mice, 2 cells. Mean and SEM are shown. No significant differences were found between groups. **e,** Proportion of neurons that fired an action potential in response to ChR2 stimulation of cortical input for each cell-type. **f,** Mean pre-light and post-light interspike interval (ISI) was measured in 4 ChAT*+* neurons and 5 Pvalb*+* neurons that were spontaneously spiking (no FoxP2+ neurons were spontaneously spiking). Example traces are shown for each cell type (timing of 470 nm 2ms light pulse is indicated by the dashed vertical line). There was no significant difference in interspike interval after light for ChAT+ neurons (Two-tailed paired t-test: t=1.303, df=3, p=0.2835). There was a significant difference in interspike interval after light for Pvalb+ neurons (Two-tailed paired t-test: t=2.910, df=4, p=0.0437).

## Discussion

The globus pallidus externa (GPe) is a heterogenous, richly connected basal ganglia nucleus that plays a pivotal role in determining basal ganglia output. Although the exact function of the GPe during behavior remains mysterious, numerous lines of evidence implicate it in both motor and reward-related aspects of goal-directed behavior,^1–3,9,10^ and GPe dysfunction has been linked to multiple neurologic and psychiatric disorders including Parkinson’s disease^1,11,41,59^, Huntington’s disease^17,18^, dystonia^14–16^, Tourette’s syndrome^19^, obsessive compulsive disorder, addiction^1,2^ and sleep disorders^13,62,63^. Canonical models of the basal ganglia describe the GPe as an internal basal ganglia nucleus, receiving input only from striatum and subthalamic nucleus and relaying that information only to other basal ganglia nuclei. However, several studies have now confirmed the existence of direct projections from the GPe to the cortex^33,34,49^ and direct cortical inputs back to the GPe^56^ in rodents as well as humans^64^, highlighting that GPe is an output nucleus of the basal ganglia that interfaces directly with cortex.

Our investigation provides a detailed profile of the anatomic, molecular and electrophysiologic properties of pallidocortical circuits. We demonstrate that pallidocortical neurons encircle the borders of GPe and maintain a topography that reflects the topography of their cortical projection targets. We show that pallidocortical neurons fall into two distinct electrophysiological clusters, segregated by intrinsic and action potential firing properties. Molecularly, pallidocortical neurons also fall into two distinct categories, *Npr3/Slc32a1-*expressing and *Nkx2-1/Chat-*expressing neurons, and these molecular markers can be harnessed to target these neurons for further study using conditional viral expression systems in transgenic mice. We also show that the GPe receives reciprocal input back from the cortex, from a broad swath of cortical regions, including non-motor and higher-order association areas. This input arises from deep layers (V/VI) of the cortex, predominantly from Rbp4*+* intratelencephalic cortical projection neurons. This contrasts with cortical inputs to the striatum, which arise from both layers II/III and layer V, and from both intratelencephalic and pyramidal tract projection neurons. Cortical input to the GPe is monosynaptic, glutamatergic, and appears to target different types of GPe neurons and circuits uniformly, including those that project to traditional targets such as the SNr as well as those that project back to the cortex. The latter projection forms a closed loop pallido-cortical-pallidal circuit. The functional impact of cortical input on GPe neural activity depends upon the receiving cell’s molecular and physiologic properties. Although inputs to ChAT+ neurons are relatively small, these easily evoke action potentials, whereas inputs to Pvalb*+* neurons do not tend to evoke new spikes but do reset the phase of spontaneously firing neurons.

The GPe is not unique as a basal ganglia nucleus in receiving direct cortical input. In addition to the known cortical projections to the primary input nucleus, the striatum, and the “hyperdirect pathway” from motor cortex to STN, we and others^65^ show that the SNr also receives direct glutamatergic input from both motor and non-motor cortical regions (Supp. Fig. 13). This suggests that fast communication between the cortex and non-striatal basal ganglia nuclei is not a rare phenomenon, but rather is a basic principle of basal ganglia structure and function. In keeping with current theories of the role of the basal ganglia in integrating both motor- and non-motor function, we show that both the GPe and the SNr receive cortical input from motor as well as non-motor associative cortex. Together these findings suggest that hyperdirect cortical-basal ganglia loops may support the integration of information about internal state and the external environment in the basal ganglia to produce rapid changes in goal-directed behavior.

There are many open questions about how reciprocal pallidocortical/corticopallidal (and corticonigral) loops contribute to the flow of neural activity through the basal ganglia. Single unit recordings of GPe neurons during stimulation of cortical input to GPe *in vivo* will allow us to build models of how cortical input is processed and filtered by different GPe cell types. In addition, to determine how this cortical input alters information flow through entire basal ganglia loops, it will be essential to simultaneously record in downstream structures such as the SNr, thalamus, striatum and cortex. Other limitations of this study include our focus on four main cortical projection targets and only two main cortical input regions. This was to provide examples of both motor and non-motor connectivity; however, there are many cortical regions that we have not explored in detail that are also likely to directly influence pallidal function, such as the orbitofrontal cortex and the retrosplenial cortex. Furthermore, it will be important to directly test the function of pallidocortical neurons and corticopallidal inputs in the regulation of behavior and particularly in goal-directed actions.

In conclusion, we confirm and extend the finding of a direct projection from GPe to the cortex, which tiles across multiple cortical regions, segregates into two distinct electrophysiological and molecular phenotypes and receives direct closed loop input back from deep layers of motor and non-motor cortex. This provides a new view of how the basal ganglia may flexibly modulate cortical function, respond to multi-modal cortical input, and produce diverse disease symptoms when disrupted. This may support identification of new therapeutic targets for neuromodulation in basal ganglia disorders.

## Acknowledgements

We thank the following funding agencies and awards for their support for this project: Conte Center for Neuroimmune Studies (grant number: GENFD0002315408), Nany Lurie Marks Foundation, The Foundation of OCD Research (BLS), American Neurological Association NRTS, Burroughs Wellcome Fund CAMS Award, NIH HRHR DP5 (grant number: 1DP5OD036140-01) (EAF). We thank the HMS Neuroimaging Facility for support for imaging and image analysis. We thank the entire Sabatini Lab for advice and support.

## Materials and Methods

### Mice

The following mouse lines were used: C57BL6/J (The Jackson Laboratory, 000664); (GENSAT, MGI: 4367067, bred in house); IT-targeting: *Tlx3-PL56-Cre* (GENSAT, MGI: 5311700, bred in house); PT-targeting: *Sim1-KJ18-Cre* (GENSAT, MGI: 4367070, bred in house), *Pvalb-IRES-Cre* (The Jackson Laboratory, 017320); Ai14 (tdTomato *Cre* reporter line, The Jackson Laboratory, 007914), *Pvalb-IRES-Cre* x *Ai14* (bred in house); *ChAT-IRES-Cre (*The Jackson Laboratory, 006410); *ChAT-IRES-Cre* x *Ai14* (bred in house); *FoxP2-IRES-Cre* (The Jackson Laboratory, 030541); *Npr3-IRES2-Cre-D* (The Jackson Laboratory, 031333). All mice were bred on a C57BL/6J genetic background and heterozygotes were used. Adult mice of either sex were used. Animals were kept on a 12:12 regular light/dark cycle under standard housing conditions. All animal care and experimental manipulations were performed in accordance with protocols approved by the Harvard Standing Committee on Animal Care, following guidelines described in the US NIH Guide for the Care and Use of Laboratory Animals.

### Intracranial injections

Mice were anesthetized with 3% isoflurane and maintained under anesthesia during surgery with 1.5% isoflurane and 80% oxygen. Using a stereotactic frame (David Kopf Instruments), the skull was exposed under aseptic conditions, a small craniotomy (around 300 μm) was drilled and cholera toxin b (CTb) or virus (non-pseudotyped rabies virus or AAV) (Table 1, Materials and Resources) was injected into the assigned brain region(s) at the coordinates listed in Table 2 (Stereotactic Injections Table). All coordinates are relative to bregma as defined in The Paxinos and Franklin Mouse Brain Altas^66^. Injection volumes and expression times are listed in Table 2 (Stereotactic Injections Table).

**Table 2.** Stereotactic injections.

| Brain region | Stereotactic coordinate (Paxinos and Franklin<sup>66</sup>) (mm) | Injection volume |
| --- | --- | --- |
| Motor cortex | Injection 1: AP +1.1, ML +/-1.3, DV 1.1, 1.35<br>Injection 2: AP +0.7, ML +/-1.2, DV 1.1, 1.35 | 150 nL at each depth |
| Sensory cortex | Injection 1: AP 0, ML +/-2.75, DV 1.5, 1.7<br>Injection 2: AP -1, ML +/-3.25, DV 1.5, 1.7 | 150 nL at each depth |
| Insular cortex | Injection 1: AP +1.5, ML +/-3.3, DV 3.4, 3.75<br>Injection 2: AP +1.1, ML +/-3.4, DV 3.2, 3.4 | 150 nL at each depth |
| Prefrontal cortex | Injection 1: AP 2.1, ML +/-0.3, DV 1.2, 1.7<br>Injection 2: AP 2.3, ML +/-0.2, DV 1.3, 2.0 | 150 nL at each depth |
| GPe | Figure 3h: AP: -0.800, ML: +/-2.25, DV: -4.000<br>Figure 4a-d: AP -0.6, ML +/-2.38, DV -4.1 | Figure 3h: 150 nL<br>Figure 4a: 75 nL |
| Striatum | Figure 4a-d: AP +0.1, ML +/-2.0, DV -3.0 | 300 nL |
| SNr | Figure 4g: AP -3.2, ML +/-1.5, DV -4.5 | 300 nL |

**Table 3.**
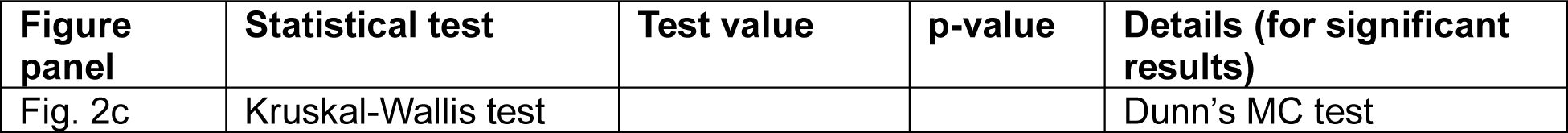
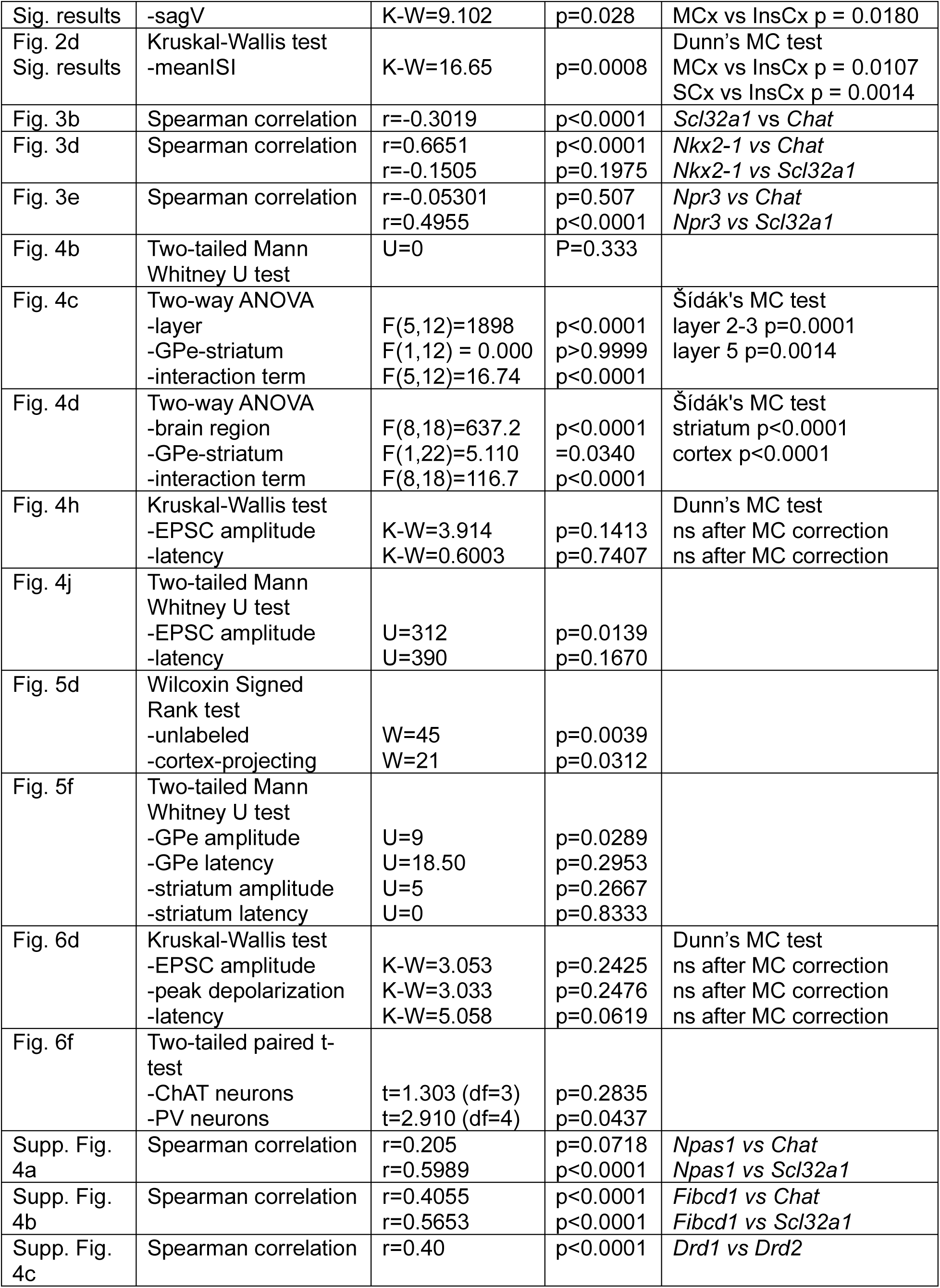
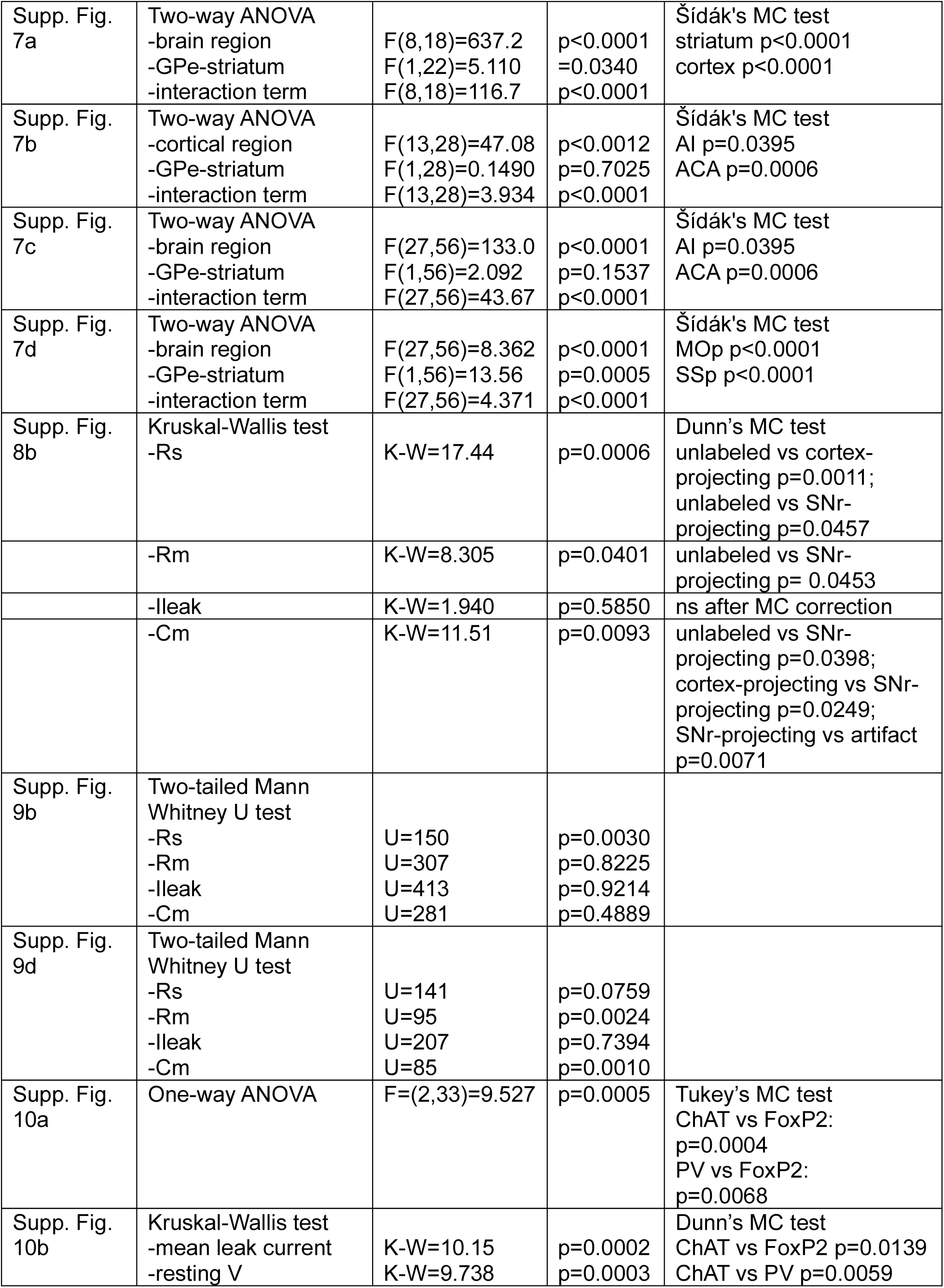
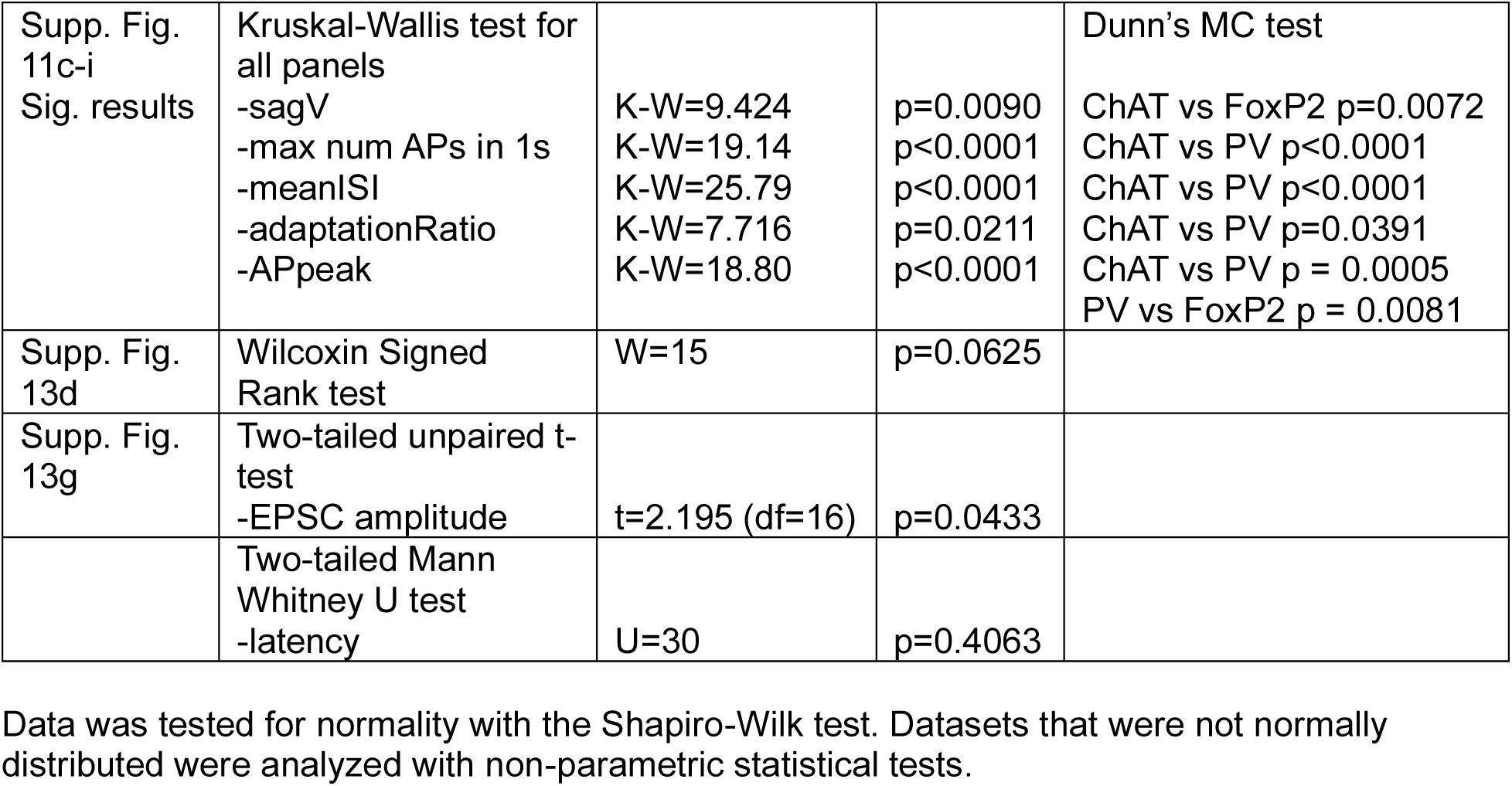
Statistical tests.

Injections were performed as previously described^67–69^. A pulled glass pipette was briefly lowered 200 µm past the target depth, then retracted and held in place at the target depth for 2 min prior to injection, then and CTb or viruses were infused at a rate of 50 nL min^−1^ using a syringe pump (Harvard Apparatus, 883015). Pipettes were slowly withdrawn at least 5 min after the end of the infusion. After injections, the wound was sutured. After surgery, mice were placed in a recovery cage with a heating pad until their activity was recovered, before being returned to their home cage. Mice were given pre- and post-operative oral carprofen (CPF, 5 mg per kg per day) analgesia and monitored daily for at least four days post-surgery. For retrograde tracer injections (rabies virus, CTb), 5-7 days were allowed for expression before experiments were performed. For *in vitro* electrophysiology experiments, ChR2 virus was allowed to express for 3-8 weeks before experiments were performed (mean expression time = 4.5 weeks, std = 1.2 weeks).

#### Rabies virus tracing injections

For transsynaptic retrograde tracing, injections were performed as described above. *ChAT-IRES-Cre* and *Npr3-IRES-Cre* mice (2-5 months) were injected with *Cre*-dependent TVA helper virus (AAV8-EF1a-FLEX-GT) into GPe, which was allowed to express for 2 weeks prior to injection of a pseudotyped rabies virus (CVS-N2c-dG-mCherry (EnvA)) into the motor or insular cortex of the same mice. Two control experiments were performed: 1) AAV8-EF1a-FLEX-GT was injected into GPe of a WT mouse to confirm that TVA expression was dependent on *Cre*-expression (Supp. Fig. 5c) 2) CVS-N2c-dG-mCherry (EnvA)) was injected into the cortex of a WT mouse (without prior TVA injection) to confirm that mCherry expression was dependent on TVA expression.

### Histology and immunohistochemistry

#### Histology for anatomical studies

Mice were deeply anesthetized with isoflurane and transcardially perfused with 5–10 ml chilled phosphate buffered saline (PBS), followed by 10–15 ml chilled 4% paraformaldehyde in PBS. Brains were dissected out and post-fixed overnight at 4°C, and transferred either to 0.1M PBS or incubated in a storing/cryoprotectant solution of 30% sucrose and 0.05% sodium azide in PBS. Brains were allowed to equilibrate for at least two days. Brains stored in PBS were sectioned into slices at 50µm thickness on a vibratome (Leica VT1000 S). Brains stored in 30% sucrose cyroprotectant solution were sectioned on a freezing microtome (Leica Biosystems, SM2010 R) before being transferred to PBS. Slices were then mounted and coverslipped with ProLong Diamond Antifade Mountant with DAPI (Thermo Fisher Scientific). Slides were imaged with an Olympus VS120 or VS200 slide scanning microscope.

#### Histology and biocytin staining for electrophysiological slices

300µm thick brain slices used in electrophysiology experiments were fixed for 24 hours in chilled 4% PFA in a 24 well plate after recordings were complete. Slices were then incubated in a storing/cryoprotectant solution of 30% sucrose and 0.05% sodium azide in PBS for at least 2 days (or until they sank). Slices were then mounted onto a block of Tissue-Tek^®^ O.C.T. Compound in a cryostat microtome (Leica Biosystems, CM1950) and re-sliced into 50 µm thick sections. The sections were then transferred to PBS for biocytin staining. For staining biocytin-filled cells with streptavidin, slices were rinsed 3 x 10 min in PBS. On the 3^rd^ wash, slices were blocked in PBS + 0.2% TritonX + 3% normal goat serum (NGS) for two hours at room temperature. After two hours of blocking, the slices were incubated in 1:1000 dilution of Streptavidin Alexa Fluor™ 647 Conjugate, 2mg/mL (Invitrogen catalog no. S32357) in blocking solution for two hours. The sections were then rinsed 3 x 10 min in PBS, before being mounted, coverslipped and imaged as described above.

### Fluorescent *in situ* mRNA hybridization (mRNA FISH) experiments

FISH experiments were performed as described previously^68,69^. Animals were deeply anesthetized with isoflurane before decapitation. Brains then were rapidly removed, embedded and frozen in tissue freezing media (Tissue-Tek^®^ O.C.T. Compound) on dry ice. 20 µm coronal sections containing GPe were prepared on a cryostat microtome (Leica Biosystems, CM1950) and mounted onto SuperFrost Plus glass slides (VWR) at 20-80 µm intervals between slices in a set. Slices were rapidly refrozen once mounted and stored at −80°C before performing *in situ* hybridization. Multiplexed fluorescent mRNA *in situ* hybridization was performed according to the ACDBio RNAscope Fluorescent Multiplex Assay protocols. Briefly, slices were fixed in 4% paraformaldehyde at 4°C for 15 min and dehydrated through washing steps in ethanol at increasing concentrations (50%, 70%, 100%) before protease digestion (Protease III, 10 min at room temperature). Probes and amplification/detection reagents were applied to the tissue sections and incubated under conditions stated in the detection protocol provided by ACDBio. Sections were counterstained using DAPI provided in the detection reagent kits and mounted in ProLong Gold mounting media (ThermoFisher Scientific P36934). Probes used are listed in Table 1 (Materials and Resources Table). Images were captured using a fluorescence microscope with structured illumination (Keyence BZ-X710) using 10X air and 60X oil immersion objectives.

### In vitro slice preparation and electrophysiology recordings with optogenetic stimulation

Brain slices were obtained from 2- to 5-month-old mice (both male and female) using standard techniques. Mice were anesthetized by isoflurane inhalation and perfused transcardially with ice-cold artificial cerebrospinal fluid (ACSF) containing (in mM) 125 NaCl, 2.5 KCl, 25 NaHCO3, 2 CaCl2, 1 MgCl2, 1.25 NaH2PO4 and 25 glucose (295 mOsm/kg). Brains were then blocked and transferred into a slicing chamber containing ice-cold ACSF. Coronal slices of GPe were cut at 300 μm thickness with a Leica VT1000s vibratome in ice-cold ACSF, transferred for 10 min to a holding chamber containing choline-based solution (consisting of (in mM): 110 choline chloride, 25 NaHCO3, 2.5 KCl, 7 MgCl2, 0.5 CaCl2, 1.25 NaH2PO4, 25 glucose, 11.6 ascorbic acid, and 3.1 pyruvic acid) at 34°C then transferred to a secondary holding chamber containing ACSF at 34°C for 10 min and subsequently maintained at room temperature (20–22°C) until use.

Electrophysiology recordings: Individual brain slices were transferred into a recording chamber, mounted on an upright microscope (Olympus BX51WI) and continuously superfused (2– 3 ml min−1) with ACSF warmed to 32–34°C by passing it through a feedback-controlled in-line heater (SH-27B; Warner Instruments). Cells were visualized through a 60X water-immersion objective with either infrared differential interference contrast optics or epifluorescence to identify cells expressing fluorophores. For whole cell voltage clamp recording, patch pipettes (2–4 MΩ) pulled from borosilicate glass (G150F-3, Warner Instruments) were filled with internal solution containing (in mM) 135 CsMeSO3, 10 HEPES, 1 EGTA, 3.3 QX-314 (Cl− salt), 4 Mg-ATP, 0.3 Na-GTP, 8 Na2-phosphocreatine (pH 7.3 adjusted with CsOH; 295 mOsm·kg−1). To record ChR2 activated EPSCs, the membrane voltages were clamped at –70 mV. Slice ChR2 fibers were stimulated with 473nm laser light, delivered at 6mW for 3ms using full-field illumination through the objective at 10 second intervals. Whole cell current clamp was used to study intrinsic properties of GPe neurons and their membrane potential responses to ChR2 stimulation. For current clamp recordings, patch pipettes were filled with internal solution containing (in mM) 135 KMeSO3, 10 HEPES, 1 EGTA, 4 Mg-ATP, 0.4 Na-GTP, 8 Na2-phosphocreatine (pH 7.3 adjusted with KOH; 295 mOsm·kg−1). At the resting membrane potential, currents were injected for 1000ms at 50pA steps from -100 to 1000pA. Optogenetic and currents injection-induced signals were acquired with a MultiClamp 700B amplifier (Molecular Devices) and digitized at 10 kHz with a National Instruments data acquisition device (NI USB-6343). Data were recorded and analyzed using a custom program written for MATLAB.

Pharmacologic manipulations: All recordings (unless specified) were performed in the presence of gabazine (10µM, Tocris) to block GABAergic neurotransmission. Additional pharmacologic manipulations were performed as described in the main text using TTX (10 µM, Tocris), 4AP (400 µM, Sigma-Aldrich), NBQX (10 µM, Tocris), CPP (µM, Sigma-Aldrich). For post-hoc histologic visualization of recorded cells, biocytin 1mg/mL (Sigma-Aldrich) was added to the internal solution. Slices were fixed in chilled 4% PFA and histology was performed as described above.

### Analysis

#### Anatomical analyses

Sagittal images of retrograde CTb tracing experiments were manually aligned to sagittal images from the Allen Brain Atlas to represent “lateral” (ML = 2.725mm), “intermediate” (ML = 2.35mm), and “medial” (ML = 1.95mm) sagittal planes. Zoomed sections of GPe for each Allen Brain Atlas Image were then used to manually annotate the location of cells expressing different CTb fluorophores in GPe, corresponding to different cortical projection targets. Co-labeling of cells with two or more CTb fluorophores was also annotated. The number of cells projecting to each cortical projection target (motor, sensory, insular, prefrontal, or colabeled cells) was quantified for each sagittal section.

#### mRNA FISH analysis

The rabies channel Keyence images were used to create a mask of rabies expression using a Otsu thresholding or a custom machine-learning algorithm in MATLAB, to automatically segment rabies-infected cells. Slices were discarded if excessive background fluorescence caused incorrect cell segmentation. RNAscope puncta quantification was performed using a custom machine learning algorithm in MATLAB (developed by the HMS Neuroimaging Facility). Puncta counts were exported as csv files for further analysis and data visualization using custom scripts in MATLAB and using GraphPad. Bootstrap analyses of the Spearman correlation coefficient for puncta counts for two different mRNA markers within individual cells was performed using 1000 bootstrap repetitions. The bootstrap analysis was repeated for the same datasets after random shuffling of the data. Poisson based minimum error thresholding was used to classify cells as “expressing” or “non-expressing” for a given mRNA marker^70,71^.

#### NeuroInfo whole brain cell quantification pipeline

Coronal images were imported to NeuroInfo software (MBF Bioscience, Williston, VT) for alignment and registration to the Allen Mouse Brain Atlas. All images were inspected and adjusted manually as necessary for correct alignment. Cells were detected automatically using the “Cell Detection Pipeline”, and all images were inspected carefully for accurate detection and edited manually as necessary to remove obvious false positive detections. The cell detection data was extracted and analyzed further using custom MATLAB scripts.

#### Electrophysiological data analysis

Electrophysiological recording traces were analyzed using custom MATLAB scripts. Quality control criteria for voltage clamp recordings were: 1) holding current Ih < -400 pA at V = -70 mV at the start of the trace (first 1000ms) and during the 1000ms prior to the light pulse (if light pulse is delivered) or at the start and end of the trace (first and last 1000ms) if no light pulse delivered (intrinsic property recordings), and 2) series resistance Rs < 30 MΩ, 3) variation in series resistance Rs < 20% over the course of all traces for that cells. Quality control criteria for current clamp recordings were: 1) resting membrane potential Vm > -90mV and < -50mV at the start and end of the trace (first and last 1000ms), 2) standard deviation of baseline membrane potential (first 1000ms) < 5mV. Cells were classified as “responders” to optogenetic stimulation if the change in membrane current after the light pulse was greater than 4 x the standard deviation of the baseline membrane current prior to the onset of the light pulse.

#### Statistical tests

Data was tested for normality with the Shapiro-Wilk test. Datasets that were not normally distributed were analyzed with non-parametric statistical tests, including the Kruskal-Wallis test (for > 2 groups), or the Mann Whitney test (unpaired data) or Wilcoxin test (paired data) for comparisons between two groups. Normally distributed data were analyzed with parametric tests including one-way ANOVA (for > 2 groups) or unpaired or paired t-tests. Mean and SEM are shown for all datasets. P values are represented by the following symbols: * for 0.01 < P < 0.05, ** for 0.001 < P < 0.01, *** for 0.0001 < P <0.001, and **** for P < 0.0001. See Table 2, Statistical Tests for further details of individual statistical comparisons.

---

**Supp. Figure 1.**
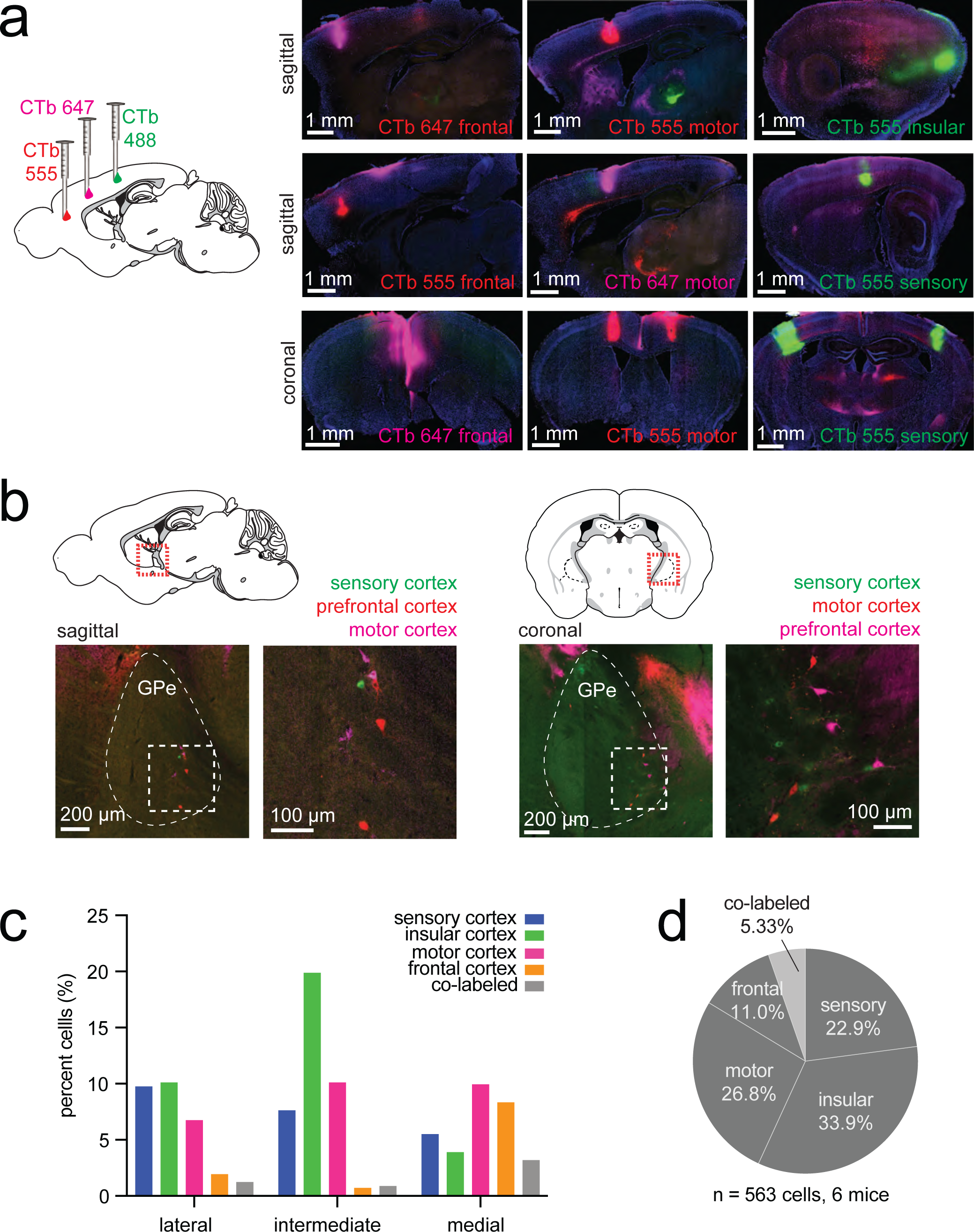
Retrograde labeling of pallidocortical neurons using triple injections of cholera toxin b (CTb). **a,** *left,* Schematic of experimental approach: CTb was injected into 3 different cortical areas in each mouse (combinations of sensory cortex, motor cortex, insular cortex, frontal cortex). *right,* Example sagittal (*top, middle rows)* and coronal (*bottom row*) images of CTb injection sites in cortex. **b,** Example sagittal and coronal images of CTb labeling in GPe. **c,** Percent of pallidocortical neurons in GPe from each cortical injection, according to sagittal plane (lateral GPe: ML 2.72 mm, intermediate GPe: ML 2.35 mm, medial GPe: ML 1.95 mm). Co-labeled cells are those in which two different CTb fluorophores were co-expressed. **d.** Percent of total number of pallidocortical neurons in GPe from each cortical injection. (total n=563 cells, across 6 mice).

**Supp. Figure 2.**
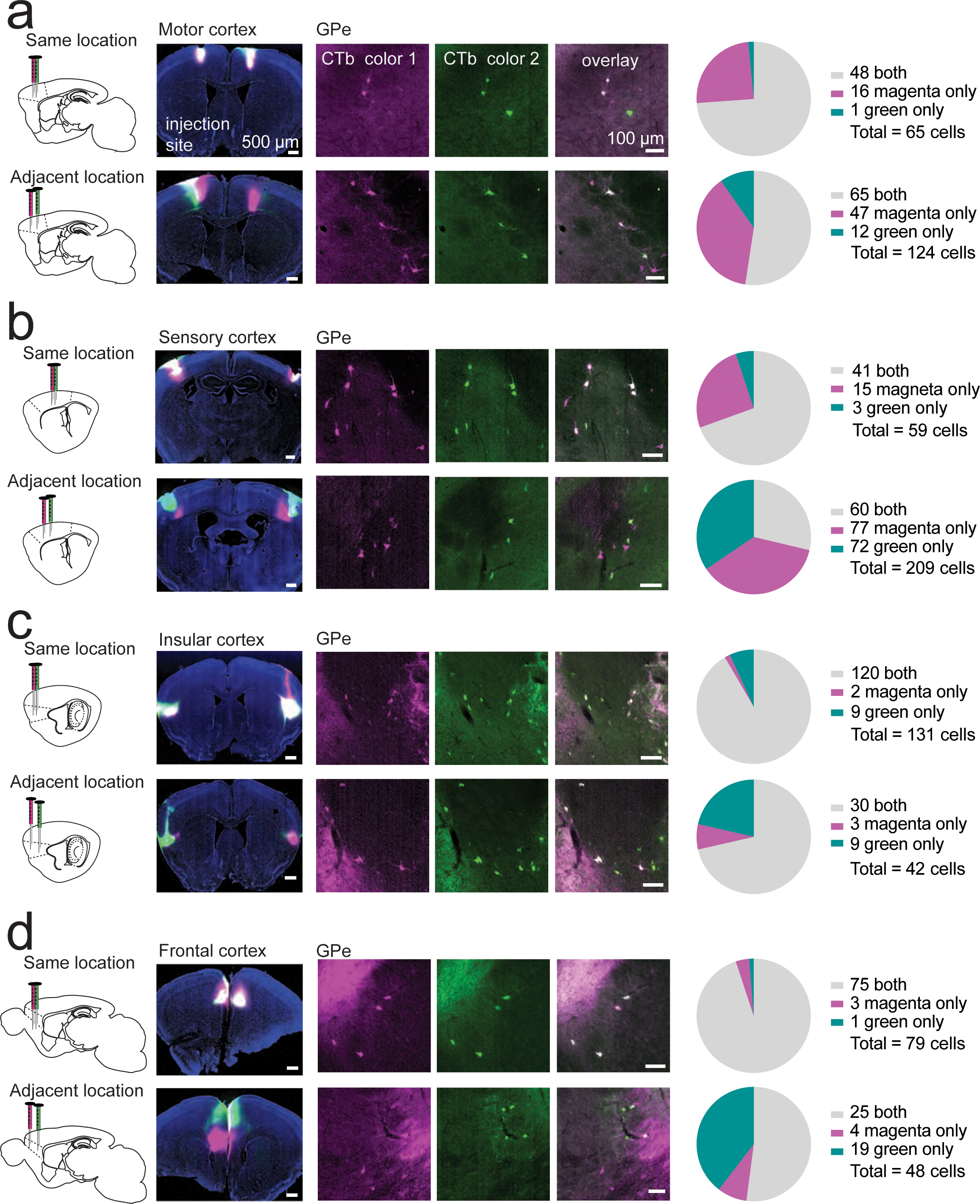
Retrograde labeling of pallidocortical neurons using double injections of CTb. **a,** *left, Schematic of* experimental approach: 2 different CTb fluorophores were injected into motor cortex, *top row,* at the same location (identical injection coordinates), or *bottom row,* at adjacent locations (slightly separated injection coordinates, please see Table 2, Stereotactic Injections) for details). *middle,* Example images of injection sites and CTb-labeled neurons in GPe (Color 1 (magenta pseudocolor)=CTb 657 or CTb 555, color 2 (green pseudocolor)=CTb 488). *right,* Quantification of co-labeling by the two different CTb fluorophores. **b,** Same as **a**, for sensory cortex. **c,** Same as **a**, for insular cortex. **d,** Same as **a**, for frontal cortex.

**Supp. Figure 3.**
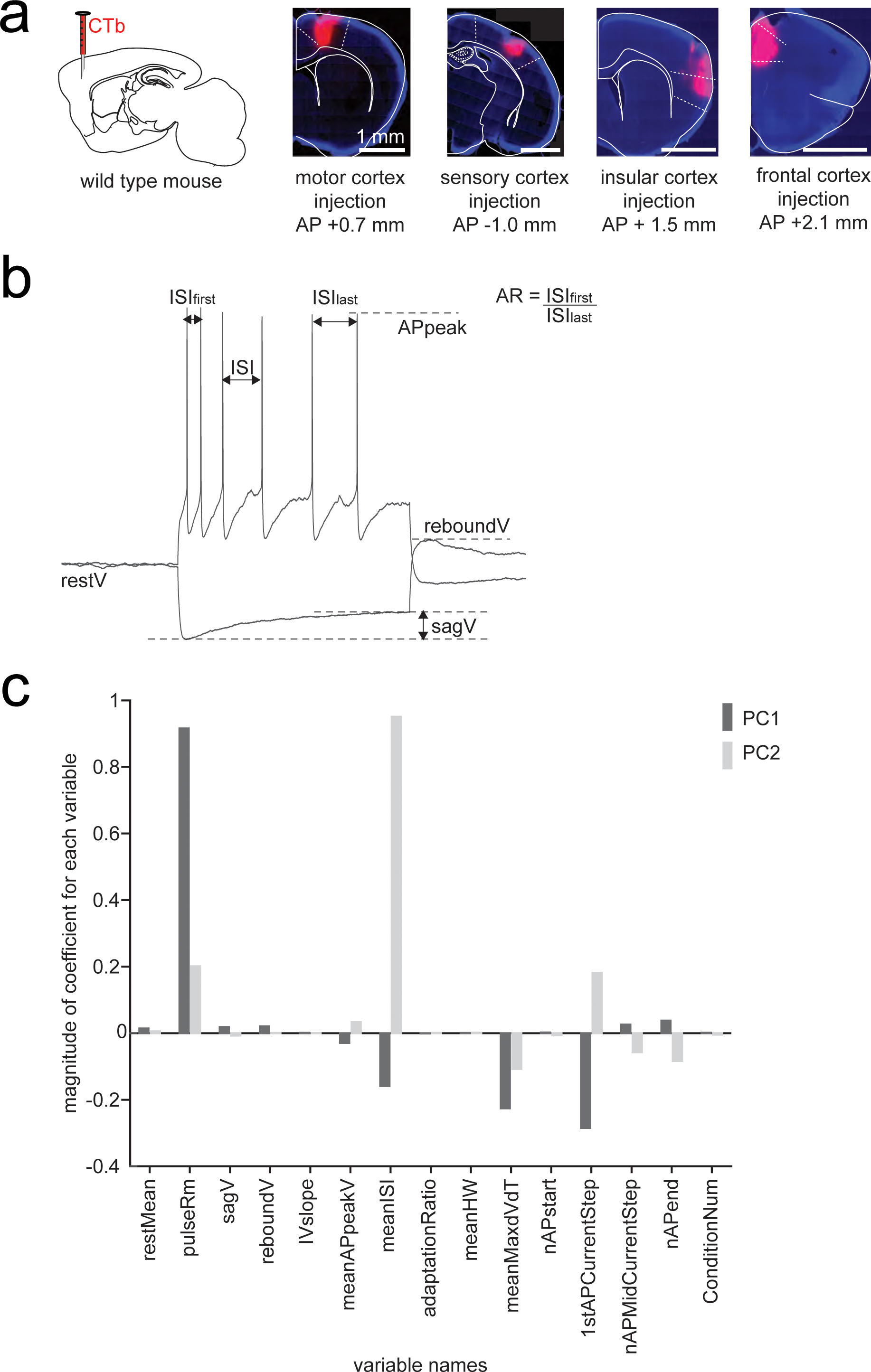
Intrinsic properties of pallidocortical neurons. **a,** *left,* Schematic of experimental approach: CTb injections into cortex. *right,* Example images of CTb fluorescence in cortex at different injection sites. **b,** Diagram of measurements for intrinsic and active membrane properties (see abbreviations below). **c,** Bar chart of variable coefficients from PCA. The bar chart shows the coefficients of the variables for the first and second principal components. The x-axis lists the variables, and the y-axis indicates the magnitude of their coefficients. The height of each bar represents the strength and direction of each variable’s contribution to the principal component; taller bars signify larger coefficients and greater influence. Positive coefficients are shown above the x-axis, while negative coefficients extend below it. *Variable name abbreviations*, restMean: resting membrane potential, pulseRm: membrane resistance, sagV: sag potential, reboundV: rebound potential, IVslope: slope of subthreshold current-voltage relationship, meanAPpeakV: mean peak voltage of all action potentials, meanISI: mean interspike interval between all action potentials, adaptationRatio: mean ratio between the first action potential pair and last action potential pair for each current step, meanHW: mean action potential half-width, meanMaxdVdT: mean maximum rate of change of membrane potential during rising phase of action potential (action potential waveform acceleration), nAPstart: number of action potentials during first current step to elicit an action potential, 1stAPCurrentStep: magnitude of current step that first elicits an action potential, nAPMidCurrentStep: number of action potentials elicited during a current step midway through the current step protocol, nAPend: number of action potentials elicited during the last current step (typically 280-400pA), conditionNum: condition number refers to the experimental group which indicates the cortical target of the recorded pallidocortical neuron.

**Supp. Figure 4.**
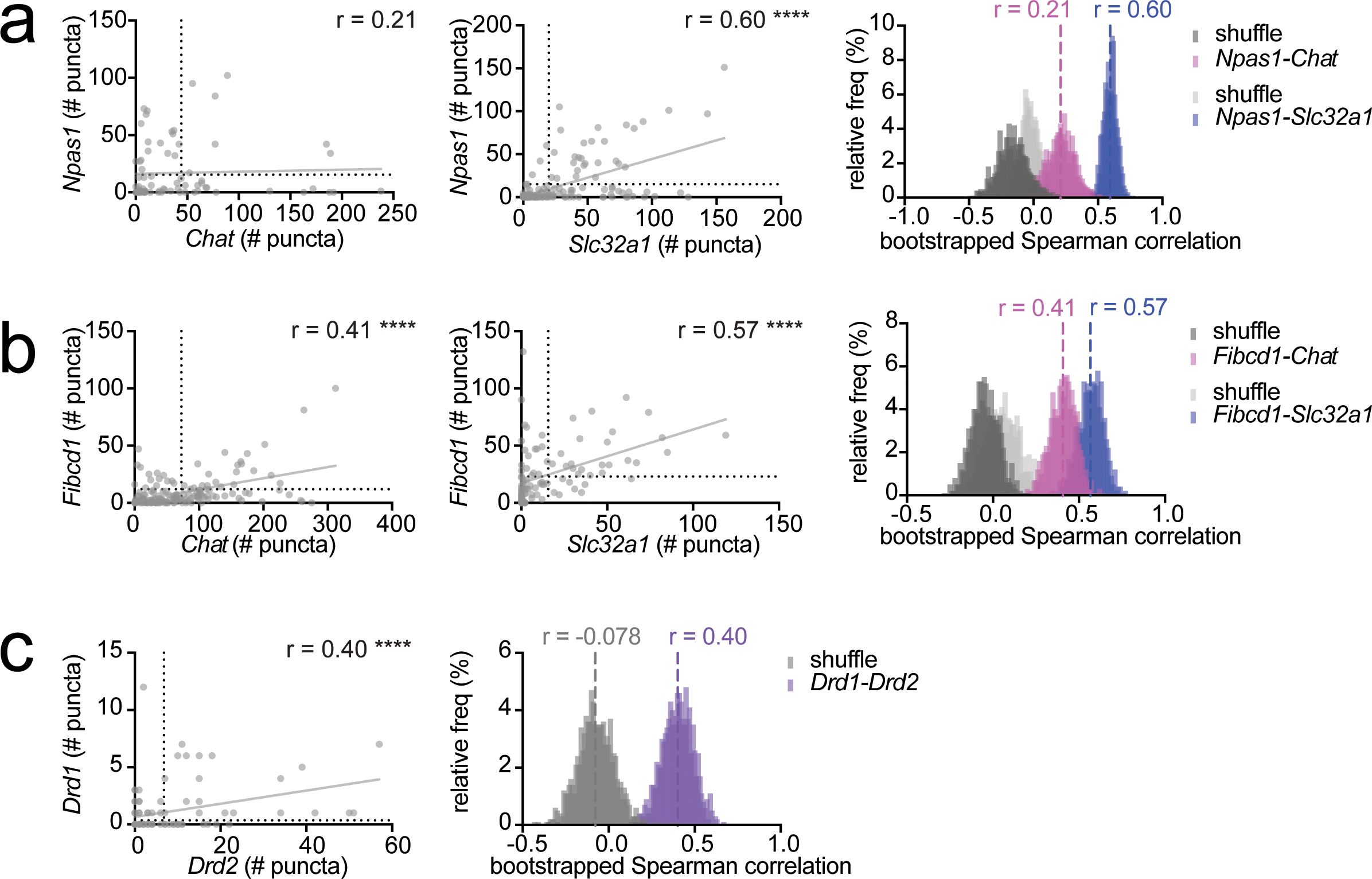
mRNA *in situ* characterization of pallidocortical neurons. **a,** Quantification of mRNA puncta for *Npas1* vs *Chat* (Spearman rho: 0.205, p=0.072, n=2 mice, 78 cells) and *Npas1* vs *Slc32a1* (Spearman rho: 0.60, p < 0.0001, n=2 mice, 197 cells*). right,* Bootstrap analysis for Spearman correlation coefficients for non-shuffled and shuffled data for *Npas1* vs *Chat* and *Npas1* vs *Slc32a1;* mean correlation coefficients for non-shuffled data are shown, along with distribution of shuffled and non-shuffled bootstrapped data. **b,** Quantification of mRNA puncta for *Fibcd1* vs *Chat* (Spearman rho: 0.41, p < 0.0001, n=2 mice, 158 cells) and *Fibcd1* vs *Slc32a1* (Spearman rho: 0.57, p < 0.0001, n=2 mice, 101 cells). Bootstrap analysis for Spearman correlation coefficients for non-shuffled and shuffled data for *Fibcd1* vs *Chat* and *Fibcd1* vs *Slc32a1;* mean correlation coefficients for non-shuffled data are shown, along with distribution of shuffled and non-shuffled bootstrapped data. **c.** Quantification of mRNA puncta for *drd1* vs *drd2* (Spearman rho: 0.40, p < 0.0001, n=1 mouse, 98 cells). Bootstrap analysis for Spearman correlation coefficients for non-shuffled and shuffled data for *drd1* vs *drd2;* mean correlation coefficients for non-shuffled data are shown, along with distribution of shuffled and non-shuffled bootstrapped data.

**Supp. Figure 5.**
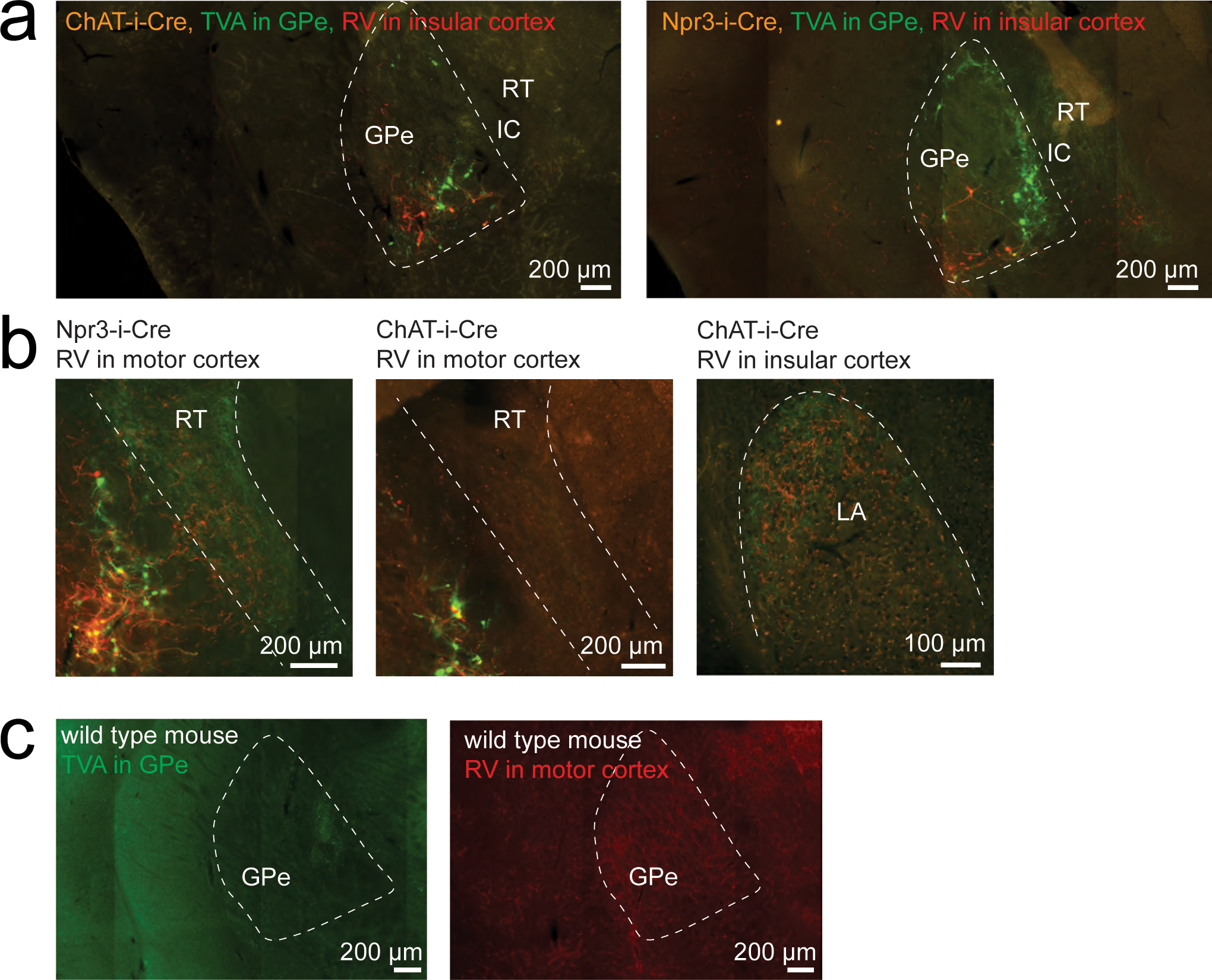
Example images of cell-type specific rabies tracing of pallidocortical neurons. **a,** Example images of ChAT+ (*left*) and Npr3+ (*right*) pallidocortical neurons that project to insular cortex. *Cre-*dependent TVA-GFP was injected into GPe, CVS-N2c-dG-mCherry (EnvA) was injected into insular cortex in either *ChAT-i-Cre* (left image) or *Npr3-i-Cre* (right image) mice. **b,** mCherry-positive rabies-expressing (CVS-N2c) fibers were observed in other brain regions, for example, in *Npr3-i-Cre* mice with motor cortex rabies virus injection, rabies-expressing fibers were observed in the reticular nucleus of the thalamus. In *ChAT-i-Cre* mice with insular cortex rabies virus injection, rabies-expressing fibers were observed in the lateral amygdala. **c,** Examples of wild-type control images: *left*, AAV-DIO-TVA-GFP was injected into the GPe of wild type mice; *right*, CVS-N2c-dG-mCherry (EnvA) was injected into motor cortex of wild type mice.

**Supp. Figure 6.**
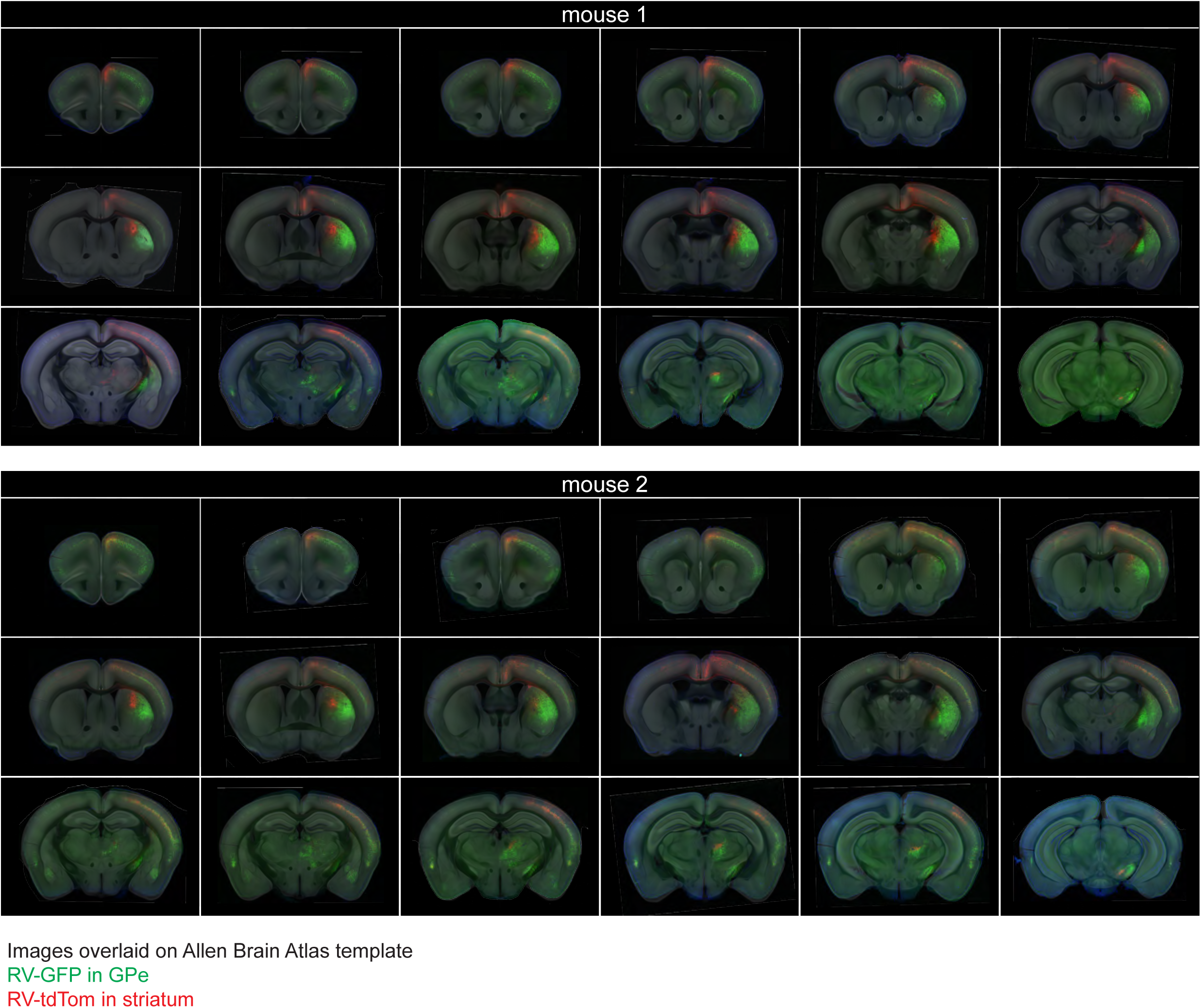
Rabies tracing of corticopallidal versus corticostriatal inputs. Example images of rabies virus expression across the brain 7 days after injection in GPe (green) or striatum (red), n=2 mice, histological images are overlaid on Allen Brain Atlas template.

**Supp. Figure 7.**
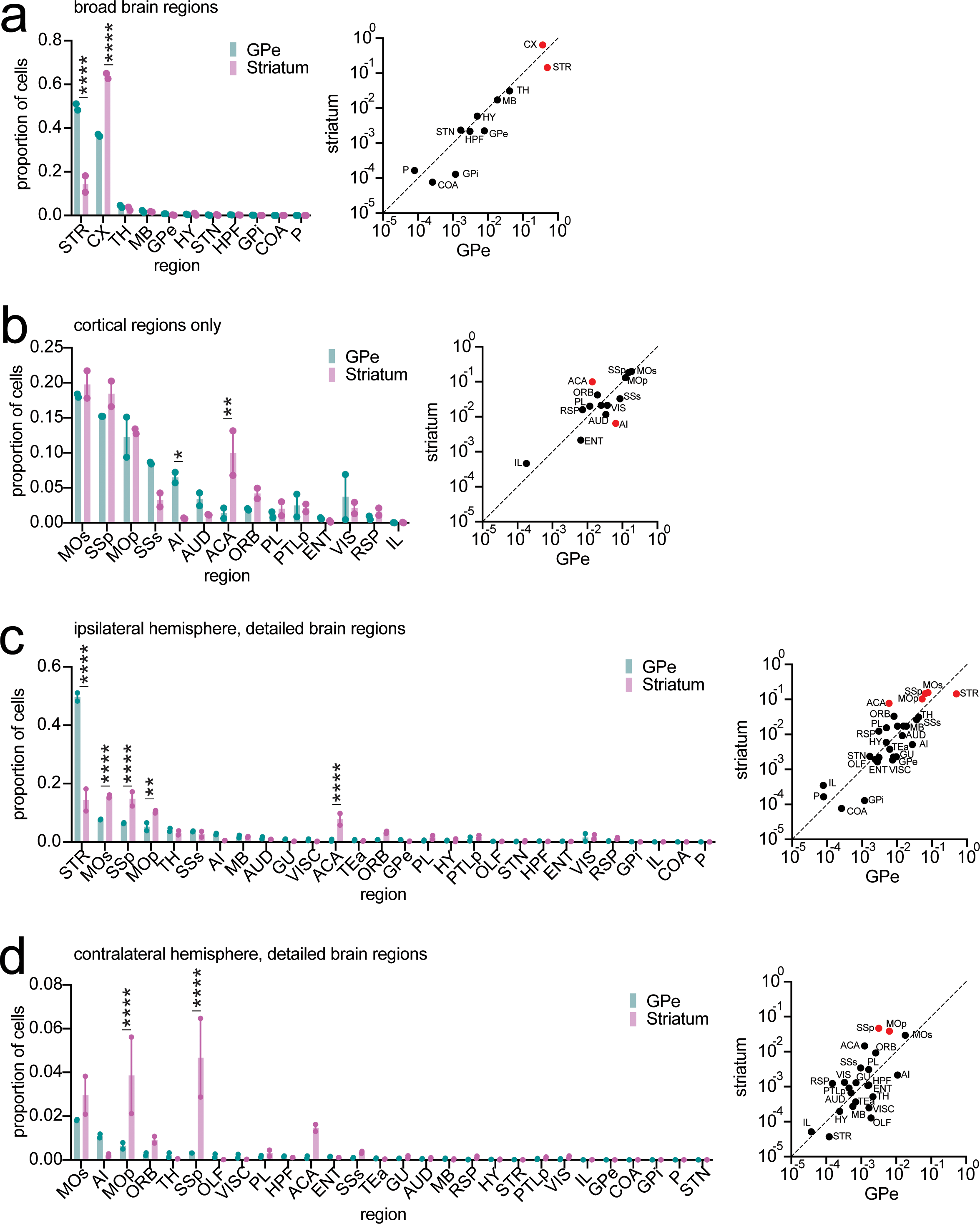
Analysis of rabies tracing of corticopallidal versus corticostriatal inputs. **a,** Proportion of ipsilateral cells innervating GPe vs striatum, stratified by broad brain regions (n=2 mice), same data as in Figure 4d. There was a significant difference between GPe and striatum for cortical input (p<0.0001) and striatal input (p<0.0001), after multiple comparison correction (two-way ANOVA with Šídák’s multiple comparison test). **b,** Proportion of all ipsilateral cortical cells innervating GPe vs striatum, stratified by cortical region (n=2 mice). There was a significant difference between GPe and striatum for agranular insular area (p=0.0395) and anterior cingulate area (p=0.0006), after multiple comparison correction (two-way ANOVA with Šídák’s multiple comparison test). **c,** Proportion of all ipsilateral cells innervating GPe vs striatum, stratified by detailed brain region (n=2 mice). There was a significant difference between GPe and striatum for the striatum, primary somatosensory cortex, secondary somatosensory cortex, secondary motor cortex and anterior cingulate cortex (p<0.0001) and primary motor cortex (p = 0.0007), after multiple comparison correction (two-way ANOVA with Šídák’s multiple comparison test). **c,** Proportion of all contralateral cells innervating GPe vs striatum, stratified by detailed brain region (n=2 mice). There was a significant difference between GPe and striatum for the primary motor and somatosensory cortices (p<0.0001) after multiple comparison correction (two-way ANOVA with Šídák’s multiple comparison test). *Key,* STR: striatum, MOp: primary motor areas, MOs: secondary motor areas, SSp: primary somatosensory areas, SSs: secondary somatosensory areas, ACA: anterior cingulate areas, ORB: orbital areas, PL: prelimbic areas, IL: infralimbic areas, AI: anterior insular area, GU: gustatory area, VISC: visceral area, AUD: auditory areas, VIS: visual areas, TEa: temporal association areas, PTLp: posterior parietal association areas, RSP: retrosplenial areas, ENT: entorhinal areas, OLF: olfactory areas, HPF: hippocampal formation, STR: striatum, GPe: globus pallidus externa, GPi: globus pallidus interna, TH: thalamus, HY: hypothalamus, STN: subthalamic nucleus, COA: cortical amygdala area, P: pons.

**Supp. Figure 8.**
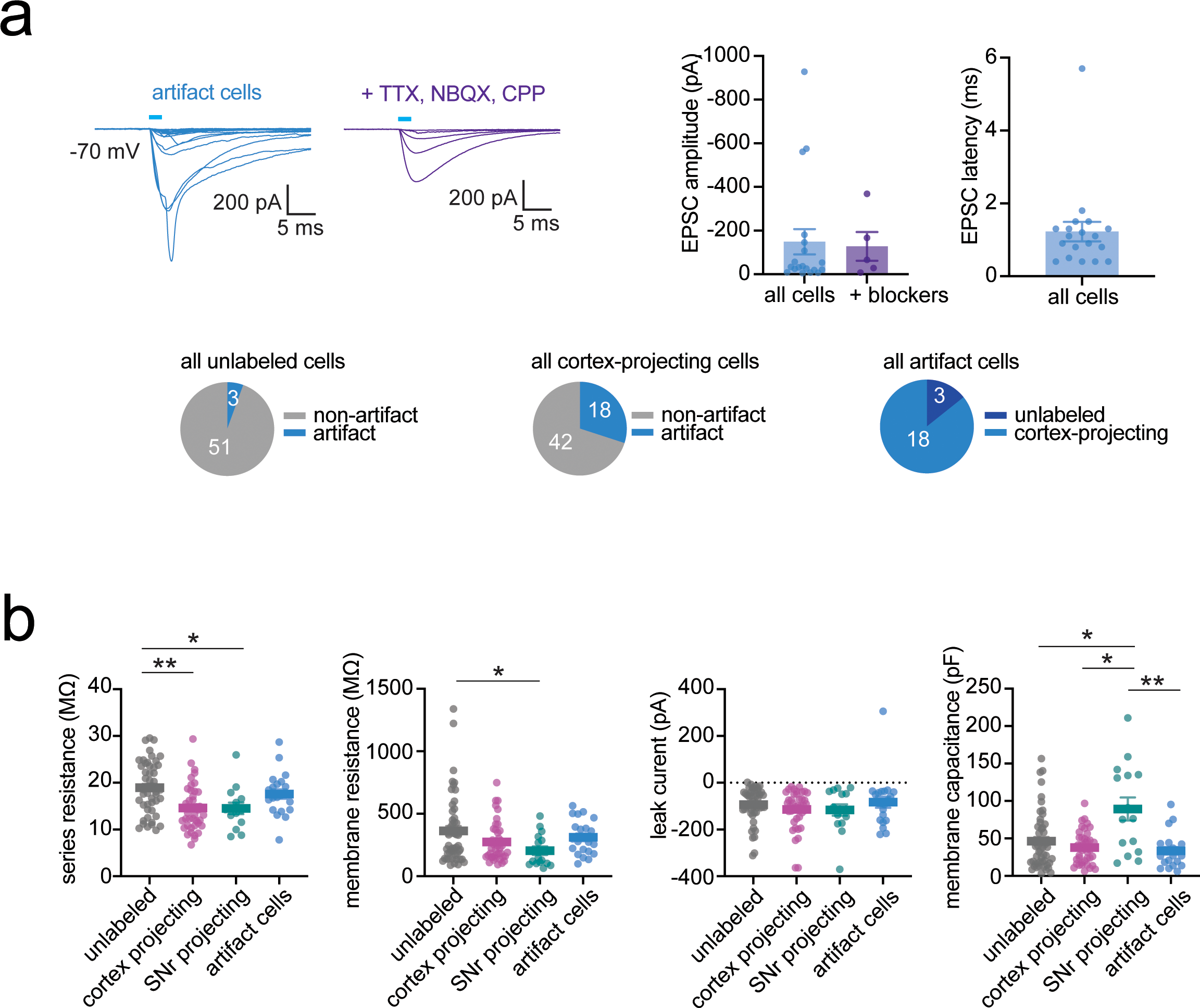
Electrophysiological characterization of cortical inputs to GPe in wild-type mice. **a,** Large rapid-onset (< 2ms from light pulse) depolarizing currents were seen in 26 cortex-projecting cells (out of a total of 69 cortex-projecting cells) and in 3 unlabeled cells (out of a total of 54 unlabeled cells). We termed these currents “artifact currents” (or photocurrents) likely related to retrograde ChR2 expression in pallidocortical neurons. *right*, Summary of EPSC amplitude and latency for all neurons with “artifact” currents. Mean and SEM are shown. **b,** Quality metrics and intrinsic properties of all neurons included in Figure 4 and Supp. Fig. 7. Mean and SEM are shown. *series resistance*, There was a significant difference in series resistance between groups (Kruskal-Wallis test K-W=17.44, p=0.0006, with the following significant results after multiple comparison correction: unlabeled vs cortex-projecting: p=0.0011, unlabeled vs SNr-projecting: p=0.0457). *membrane resistance*, There was a significant difference in membrane resistance between groups (Kruskal-Wallis test K-W=8.305, p=0.0401, with the following significant results after multiple comparison correction: unlabeled vs SNr-projecting: p=0.0453). *membrane leak current*, There was no significant difference in membrane leak current between groups (K-W=1.940, p=0.5850). *membrane capacitance*, There was a significant difference in membrane capacitance between groups (K-W=11.51, p=0.0093, with the following significant results after multiple comparison correction: unlabeled vs SNr-projecting: p= 0.0398, cortex-projecting vs SNr-projecting: p=0.0249, SNr-projecting vs artifact: p=0.0071).

**Supp. Figure 9.**
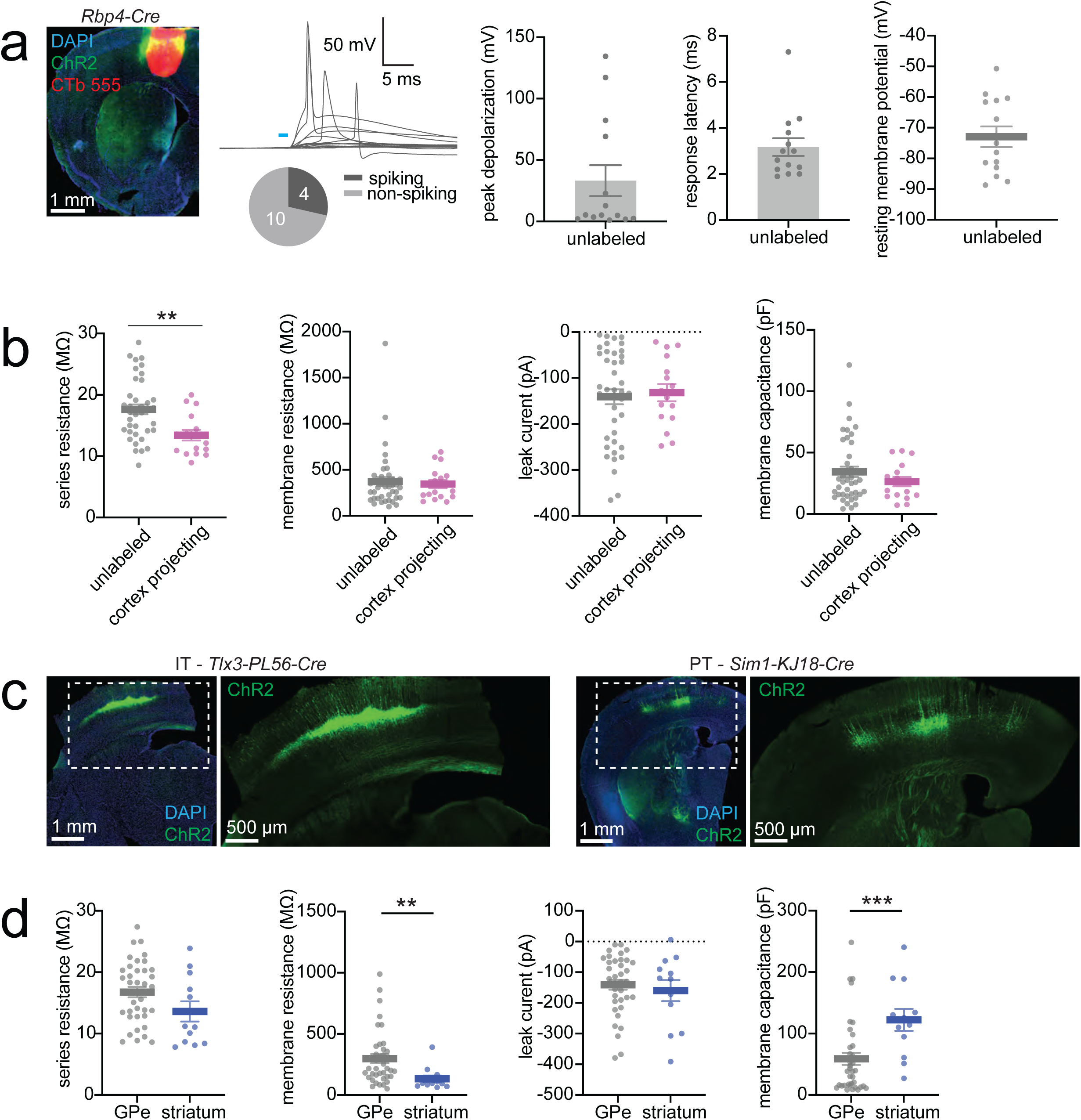
Current clamp responses of GPe neurons to *Rbp4+* cortical neuron input. **a,** *left,* Example image of motor cortex ChR2 and CTb 555 expression in *Rbp4-Cre* mouse. *middle,* Unlabeled GPe neuron current clamp responses to ChR2 stimulation of Rbp4*+* cortical axons. *right,* Peak depolarization, response latency and resting membrane potential of all recorded neurons are shown (mean and SEM are shown, n=14 cells, 7 mice). **b,** Quality metrics and intrinsic properties of all neurons from *Rbp4-Cre* experiments (Fig. 5), mean and SEM are shown. Unlabeled neurons: n=40 cells, 11 mice, cortex-projecting neurons: n=16 cells, 8 mice. There was a significant difference in series resistance between unlabeled and cortex-projecting neurons (Mann Whitney test U=150, p=0.0030). There was no significant difference in membrane resistance, leak current or capacitance between the neuron types. **c,** Example images of ChR2 expression in *Tlx3-PL56-Cre* (IT neurons, with cross-callosal intratelencephalic fibers) or *Sim1-KJ18-Cre* (PT neurons, with descending pyramidal tract fibers). **d,** Quality metrics and intrinsic properties of all neurons from *Tlx3-PL56-Cre* and *Sim1-KJ18-Cre* experiments (Fig. 5), mean and SEM are shown. GPe neurons: n=37 cells, 4 mice; striatal neurons: n=12 neurons, 4 mice. There was a significant difference in membrane resistance and membrane capacitance between GPe and striatal neurons. Membrane resistance: Mann Whitney test: U=95, p=0.0024. Membrane capacitance: U=85, p=0.0010. There was no significant difference in series resistance or leak current between the neuron types.

**Supp. Figure 10.**
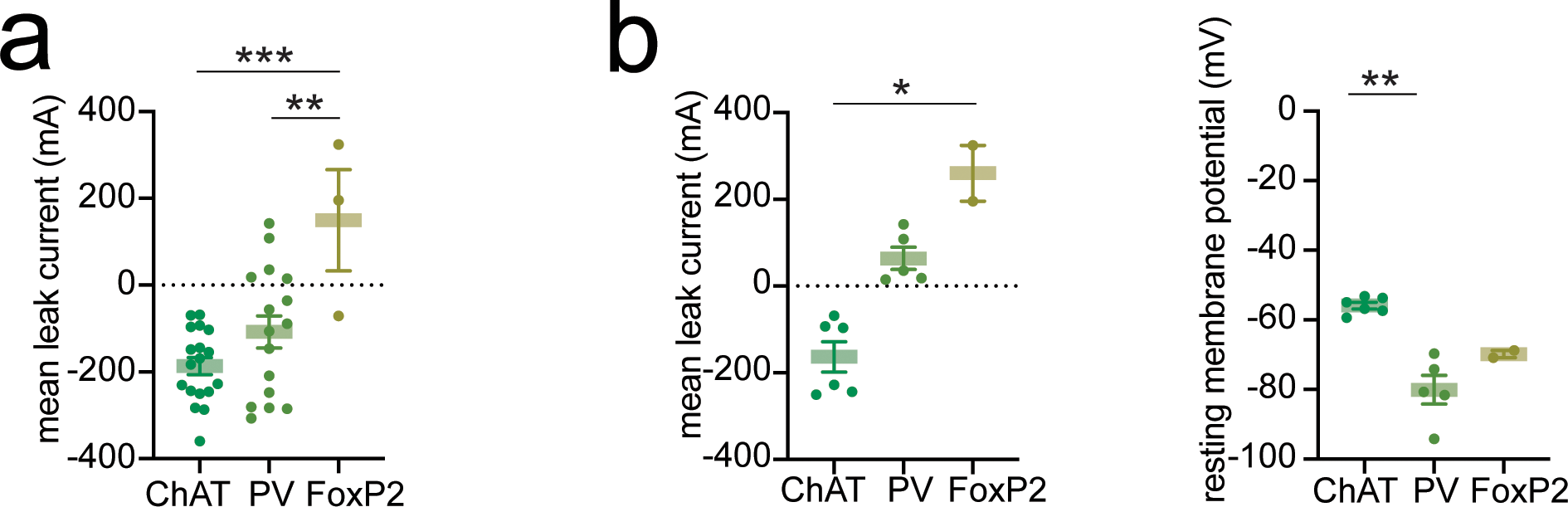
Responses to light stimulation in neurons that are spontaneously spiking. **a,** mean leak current for all neurons shown in traces in Fig. 6b, mean and SEM are shown. ChAT+ n=2 mice, 18 cells. Pvalb+ n=2 mice, 16 cells, FoxP2+ n=2 mice, 2 cells. There was a significant difference in leak current between cell types (one-way ANOVA: F (2, 33)=9.527, p=0.0005, multiple comparison correction: ChAT+ vs FoxP2+ p=0.0004; Pvalb+ vs FoxP2*+* p=0.0068. **b,** Leak current and resting membrane potential for neurons used for quantifying paired voltage clamp and current clamp responses to light stimulation in Fig. 6d, mean and SEM are shown. ChAT+ n=2 mice, 6 cells. Pvalb+ n=2 mice, 5 cells, FoxP2+ n=2 mice, 2 cells. There was a significant difference in leak current between cell types (Kruskal-Wallis test: K-W=10.15, p=0.0002, multiple comparison correction: ChAT+ vs FoxP2+ p=0.0139). There was a significant difference in resting membrane potential between cell types (K-W=9.738, p=0.0003, multiple comparison correction: ChAT+ vs Pvalb+: p=0.0059).

**Supp. Figure 11.**
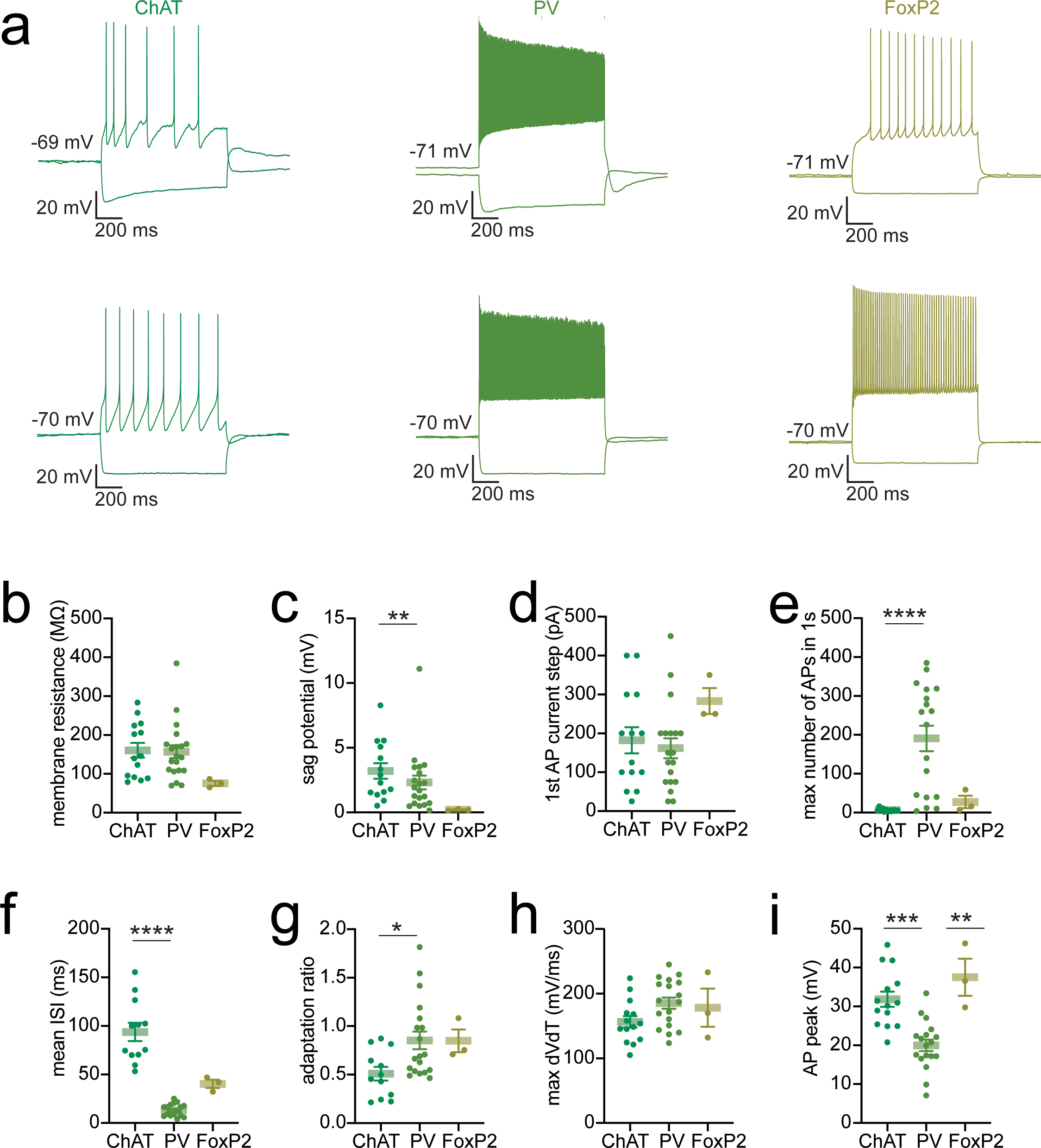
Intrinsic properties of different GPe cell types. **a,** Example current clamp recordings from different GPe cell types. A hyperpolarizing response and a supra-threshold depolarizing response is shown for each cell. **b.** Quantification of intrinsic properties for different GPe cell types. Mean and SEM are shown. ChAT+ n=14 neurons, 2 mice, Pvalb+ n=20 neurons, 2 mice, FoxP2+ n= 3 neurons, 2 mice. There was no significant difference in membrane resistance, current step required to generate a first action potential (1st AP current step), or mean action potential waveform acceleration (mean max dVdT), after multiple comparison correction. There was a significant difference between groups for the following features: sag potential (Kruskal-Wallis test: K-W=9.424, p=0.0090, significant results after multiple comparisons correction: ChAT+ vs FoxP2+ p=0.0072); max firing frequency (max number of APs in 1 s) (K-W=19.14, p < 0.0001, ChAT+ vs Pvalb+ p < 0.0001); mean inter-spike interval (ISI) (K-W=25.79, p < 0.0001, ChAT+ vs Pvalb+ p < 0.0001); adaptation ratio (K-W=7.716, p=0.0211, ChAT+ vs Pvalb+ 0.039), and peak action potential depolarization (AP peak) (K-W=18.80, p < 0.0001, ChAT+ vs Pvalb+ p=0.0005; Pvalb+ vs FoxP2+ p=0.0081).

**Supp. Figure 12.**
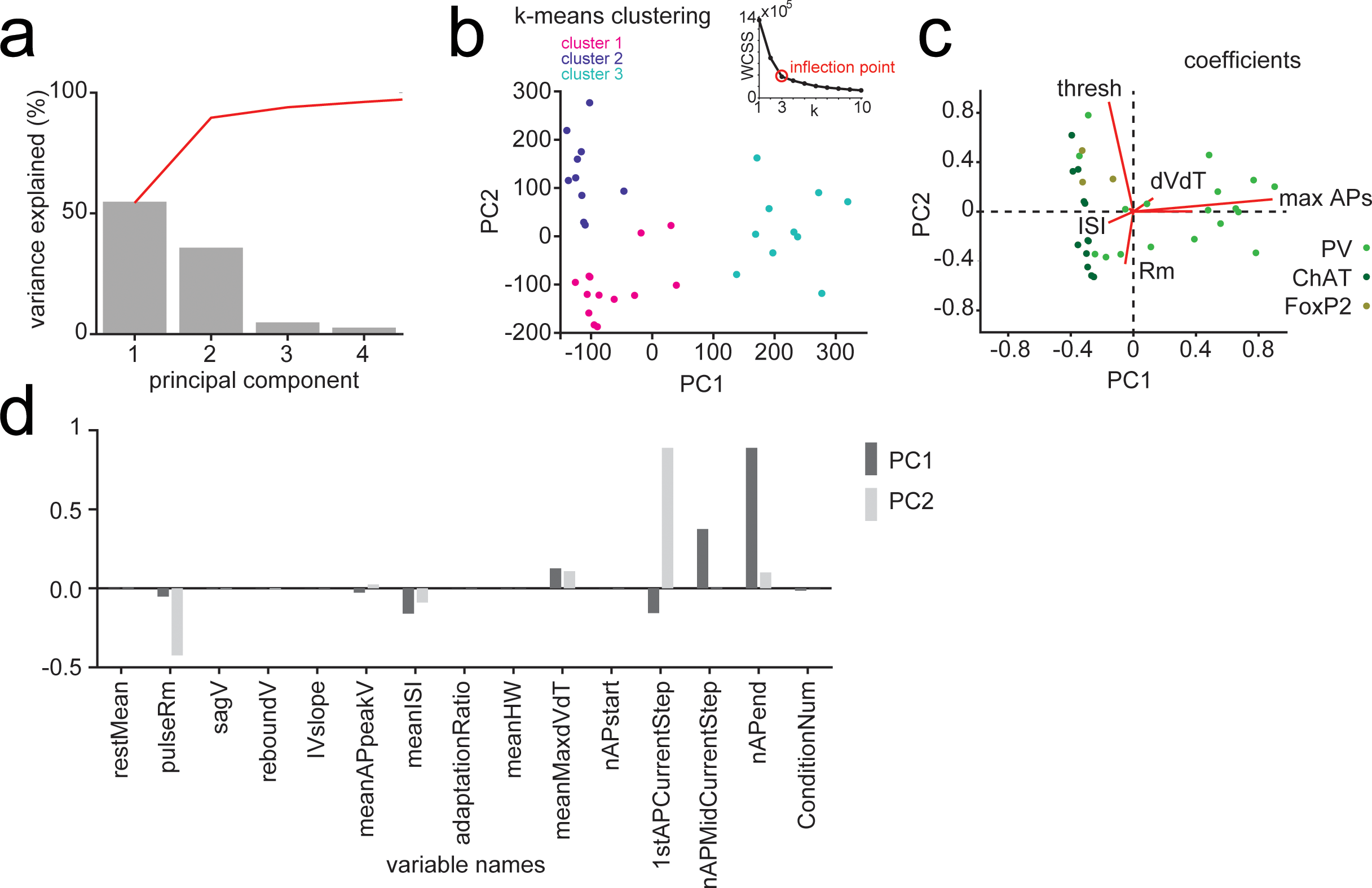
Principal components analysis for intrinsic properties of different GPe cell types. **a,** Scree plot results. Percent variance in the data explained by the first four principal components. **b,** K-means clustering results with 3 clusters. *inset:* Elbow plot of Within-Cluster Sum of Squares (WCSS) for different values of k. The maximum of the ratio of the second derivative/first derivative was identify the optimal value for k (the inflection point of the curve). Data points are color-coded by cluster assignment. The plot illustrates the distribution of data across clusters, n=6 mice, 33 cells. **C,** Biplot of the first two principal components (PC1 and PC2) showing variable coefficients and scores. Data points represent the scores for each cell on PC1 and PC2 and are color-coded according to their genetic identity (ChAT+, Pvalb+, FoxP2+). Red lines indicate the direction and magnitude of variable coefficients, with length proportional to the contribution of each variable to the principal components. **d,** Bar chart of variable coefficients from PCA. The bar chart shows the coefficients of the variables for the first and second principal components. The x-axis lists the variables, and the y-axis indicates the magnitude of their coefficients. The height of each bar represents the strength and direction of each variable’s contribution to the principal component; taller bars signify larger coefficients and greater influence. Positive coefficients are shown above the x-axis, while negative coefficients extend below it. *Variable name abbreviations*, restMean: resting membrane potential, pulseRm: membrane resistance, sagV: sag potential, reboundV: rebound potential, IVslope: slope of subthreshold current-voltage relationship, meanAPpeakV: mean peak voltage of all action potentials, meanISI: mean interspike interval between all action potentials, adaptationRatio: mean ratio between the first action potential pair and last action potential pair for each current step, meanHW=mean action potential half-width, meanMaxdVdT: mean maximum rate of change of membrane potential during rising phase of action potential (action potential waveform acceleration), nAPstart: number of action potentials during first current step to elicit an action potential, 1stAPCurrentStep: magnitude of current step that first elicits an action potential, nAPMidCurrentStep: number of action potentials elicited during a current step midway through the current step protocol, nAPend: number of action potentials elicited during the last current step (typically 280-400pA), conditionNum: condition number refers to the cell-type identity.

**Supp Figure 13.**
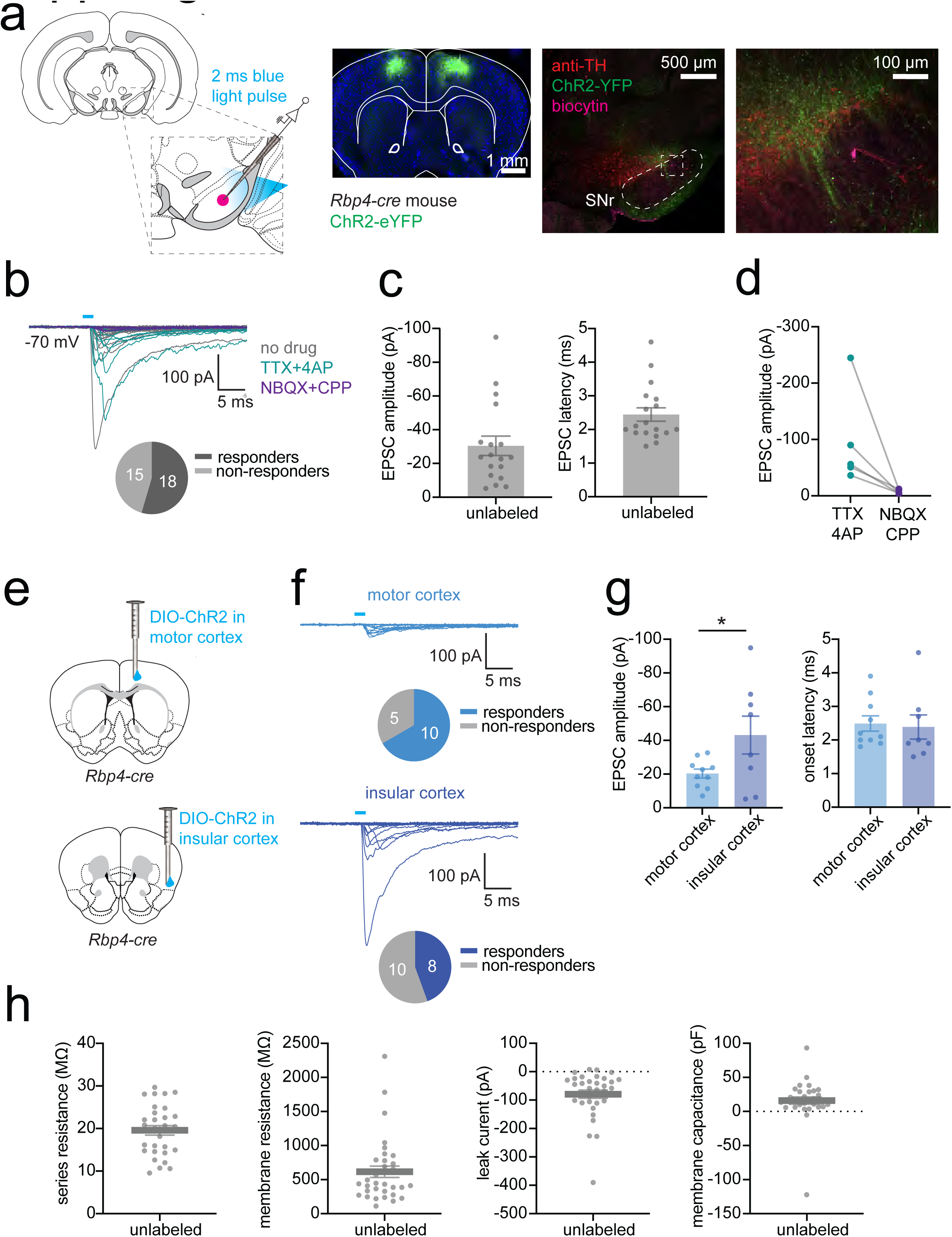
Cortical inputs to neurons in the substantia nigra pars reticulata (SNr). **a,** *left,* Schematic of experimental approach: *Cre-*dependent ChR2 was injected into the cortex of *Rbp4-Cre* mice. Neurons in the SNr were patched during stimulation of ChR2 with 2 ms pulses of 470 nm light. *right,* Example histological images of the cortical injection site and patched neurons in the SNr are shown. Anti-tyrosine hydroxylase (TH) stain is used to indicate the location of dopamine neurons in the substantia nigra pars compacta (SNc) which is dorsal to the SNr. **b,** Voltage clamp responses of SNr neurons to stimulation of cortical axons in the SNr in the absence of any pharmacologic agents, and in the presence of TTX+4AP (to isolate monosynaptic inputs) and glutamatergic receptor blockers (NBQX and CPP) to block glutamatergic input. Proportion of neurons responding to cortical input is shown below the traces. **c,** Quantification of EPSC amplitude and EPSC latency in all responding cells. Mean and SEM are shown, n=18 neurons, 5 mice. **d,** EPSC amplitude in the absence and presence of glutamatergic blockers, n=5 neuron pairs from 4 mice (Wilcoxin matched-pairs signed rank test: W=15.00, p=0.0625). **e,** ChR2 was injected either into motor cortex or insular cortex to compare inputs to SNr from two different cortical sites. **f,** Voltage clamp responses to blue light stimulation of motor cortex and insular cortex inputs. Proportion of responding neurons are shown below the traces. **g**, Quantification of EPSC amplitude and latency. Mean and SEM are shown. Motor cortex n=15 cells, 2 mice, insular cortex n=18 cells, 3 mice. There was a significant difference between motor cortex and insular cortex EPSC amplitude (two-tailed unpaired t-test: t=2.195, df=16, p=0.0433), but not EPSC latency (Mann Whitney Test: U=30, p=0.4063). **h,** Summary of intrinsic properties of all SNr neurons.

